# Centromere-proximal crossovers disrupt proper homologous chromosome disjunction during meiosis

**DOI:** 10.1101/2024.11.18.624114

**Authors:** Sucharita Sen, Sneha Sarkar, Gopika Menon, Mridula Nambiar

**Affiliations:** Department of Biology, Indian Institute of Science Education and Research, Pune 411008, Maharashtra, India

**Keywords:** Meiotic recombination, Pericentric heterochromatin, Cohesins, Chromosomal mis-segregation, Aneuploidy

## Abstract

Centromere-proximal crossovers (C-COs) are repressed during meiosis across all species. Moreover, aberrant C-COs are strongly correlated with meiotic aneuploidy such as in Down syndrome. Despite decades of work in understanding C-CO repression, the molecular basis of how they cause chromosomal mis-segregation is unclear. Here, we show that increased C-COs result in mis-segregation of homologs during meiosis I in *Schizosaccharomyces pombe*. C-COs cause either nondisjunction events where the entire bivalent moves into the same nucleus at meiosis I or result in biorientation of sister chromatids leading to their premature separation. Since meiosis I segregation appears normal in pericentric cohesion deficient mutants, we rule out centromeric cohesion disruption as the primary driver of segregation defects due to C-COs, as suggested in some other species. In contrast, reduced pericentric cohesion alleviates the meiosis I nondisjunction events, thereby supporting the previously suggested “entanglement model” that proposes physical entwining of the bivalent due to retention of sister-chromatid cohesion at centromeres, a hallmark of anaphase I. This alteration also uncovers biorientation of sister-chromatids in meiosis I suggesting mono-orientation disruption as a parallel way to promote mis-segregation in the presence of C-COs. These molecular insights will improve our understanding of infertility and aneuploidy-associated developmental disorders in humans.

## Introduction

Meiosis involves segregation of homologous chromosomes during the first round of division, while ensuring that the sister-chromatids stay held together, followed by meiosis II, where sister chromatids split. Disruptions in these mechanisms result in unequal distribution of chromosomes in the four gametes at the end of meiosis, which can severely affect their viability and lead to congenital developmental disorders such as Down syndrome characterized by presence of extra chromosomes^1^. Genetic recombination is critical during meiosis and allows shuffling of genes between the two parental chromosomes resulting in enhanced genetic diversity in the progeny. Recombination is mediated by homology-directed DNA repair in response to programmed double-strand breaks (DSBs) generated by the evolutionarily conserved Spo11, in complex with other associated proteins^2^. The recombination intermediates can be resolved to either result in a crossover or non-crossover event, the former resulting in a matured chiasma that can be seen as physical connections between the homologs in meiosis I (MI)^3^.

Apart from generating diversity, crossovers contribute towards facilitating proper separation of homologs during MI along with cohesin complexes that hold the sister-chromatids together after replication^4^. Systematic removal of these cohesin complexes helps regulate meiotic chromosome segregation – the cohesins in the arms are removed at anaphase I, but those at the pericentric regions are protected till meiosis II (MII) by Shugoshin (Sgo1), to facilitate proper segregation, similar to mitosis^5,6^. The distribution of crossovers is not uniform across the genome and are deleterious, if they occur within or near centromeres^7^. Studies in many diverse species done till date demonstrate the near absence of crossovers at the centromeres as well as the pericentric regions and a distance-dependent reduced frequency in the flanking euchromatin regions, better known as the centromere effect^8^.

The mechanisms by which this repression is mediated is found to vary among species, owing to differences in complexities in centromere organisation and the associated factors. Notably, stark differences in the repression mechanisms exist between the two highly diverged yeast species, *Saccharomyces cerevisiae* and *Schizosaccharomyces pombe*. At budding yeast point centromeres, this repression is mediated mainly by the inner-kinetochore Ctf19 complex that helps deposit cohesins in the flanking pericentric regions and consequently favor inter-sister DSB repair over the expected inter-homolog repair, thereby preventing formation of crossovers^9,10^. In contrast, at fission yeast centromeres, the centromere-specific cohesin complex containing the meiotic kleisin subunit Rec8 and the HEAT repeat associated with kleisin (HAWK) subunit Psc3, mediates the repression of meiotic crossovers at the pericentric regions flanking the centromeres^11^. Exclusion of the chromosome arms-specific pro-recombinogenic cohesin complex consisting of Rec11 (meiotic paralog of Psc3) from the centromeres, prevents activation of the DSB-inducing Rec12 (Spo11 ortholog) complex from initiating DSB formation and further homology-directed repair. Forceful targeting of the Rec12 complex-activating protein Rec10 in an otherwise wild type genetic background or targeting the Rec8-Rec11 cohesin complex to the pericentric regions in the absence of Psc3, dramatically increased DSBs and consequently centromeric crossovers (C-COs)^11^. Hence, this repression of recombination is extremely well-regulated and its absence has been positively correlated to the occurrence of trisomies and aneuploid foetuses resulting in miscarriages in humans^12^.

Increased chromosomal mis-segregation events that are seen in older oocytes from many species such as flies, mice and humans can be mainly attributed to improper recombination as well as the absence of cohesin turnover that results in weakened centromeric and sister-chromatid cohesion^13^. The former includes both absence of chromosomal crossover events or presence of aberrant centromere-proximal crossovers. Multiple linkage studies in human trisomy have found a strong correlation between centromere-proximal crossovers and the occurrence of non-disjunction (NDJ) events^14–16^. A study in budding yeast *Saccharomyces cerevisiae* showed elevated premature separation of sister chromatids (PSSC) in MI in presence of C-COs^17,18^. In contrast, spontaneous X chromosome NDJs in *Drosophila* as well as human trisomy 21 cases showed that centromere-proximal exchanges were associated primarily with MII NDJ^16,19,20^. Increased MI NDJ in cohesion-intact centromeres has been reported in human oocytes from younger females, which progressively shifts to a PSSC defect with increasing age and consequent weakening of centromeric cohesion^21^. However, despite being an age-old problem, the mechanistic basis of how C-COs result in chromosomal mis-segregation is not completely understood in any species. This question is particularly of interest as crossovers occur amply in the chromosomal arms to facilitate genetic diversity as well as promote faithful MI homolog segregation, but cause mis-segregation when present at the centromeres.

Here, we employed *S. pombe* with activated C-COs to determine the source of segregation defects during meiosis. We find that the segregation errors arise primarily due to non-disjunction of the homologs at MI and additionally likely via disruption in mono-orientation leading to premature separation of sister chromatids. The origin of these MI errors cannot be attributed to reduced centromeric cohesins, disrupted pericentric heterochromatin or decreased levels of arm crossovers. We provide evidence for a critical role of the residual protected pericentric cohesin population at the end of MI, to contribute towards non-disjunction of homologs in the presence of C-COs, which is in contrast to its fundamental role in promoting accurate segregation in MII. Hence, the entanglement theory of homologs, due to lack of resolution of chiasma, explains the segregation defects mediated by C-COs. Removal of cohesins in the presence of C-COs elevates the potential for additional MII NDJ of sister chromatids, in the nuclei that already have undergone NDJ at meiosis I. Since, *S. pombe* pericentric and centromeric regions are excellent experimental substitutes for mammalian centromeres, our study will be relevant in better understanding the defects arising during oogenesis and its association with genetic disorders. We propose that presence of C-COs predisposes the homolog non-disjunction to occur at meiosis I, which with increasing age in humans, may lead to premature separation of sister chromatids and exhibit apparent meiosis II non-disjunction phenotype in the resulting defective egg, as suggested by previous studies^15,20–22^.

## Results

### Different types of mis-segregation defects can be captured based on the source of the erroneous step during meiosis

In order to quantifiably observe and categorise chromosome segregation defects during the two stages of meiosis, we employed the previously developed fluorescence-based tetrad assay in *S. pombe*^23^. The fluorophore gene is inserted close to *cen1* (centromere of chromosome 1) under the effect of a spore-autonomous promoter. Upon completion of meiosis, the four spores within the ascus are arranged in an ordered manner, such that spores on each half of the ascus are formed from the same MI nucleus and are hence like-coloured (Fig. 1A, tetrad 1; 1B and Suppl. Fig. 1A). Any deviation from such a visual profile would mean deviation from a normal segregation pattern. We worked out additional possible categories of segregation error profiles that weren’t discussed previously, and split the errors originating in MI and MII into subtypes.

**Figure 1.**
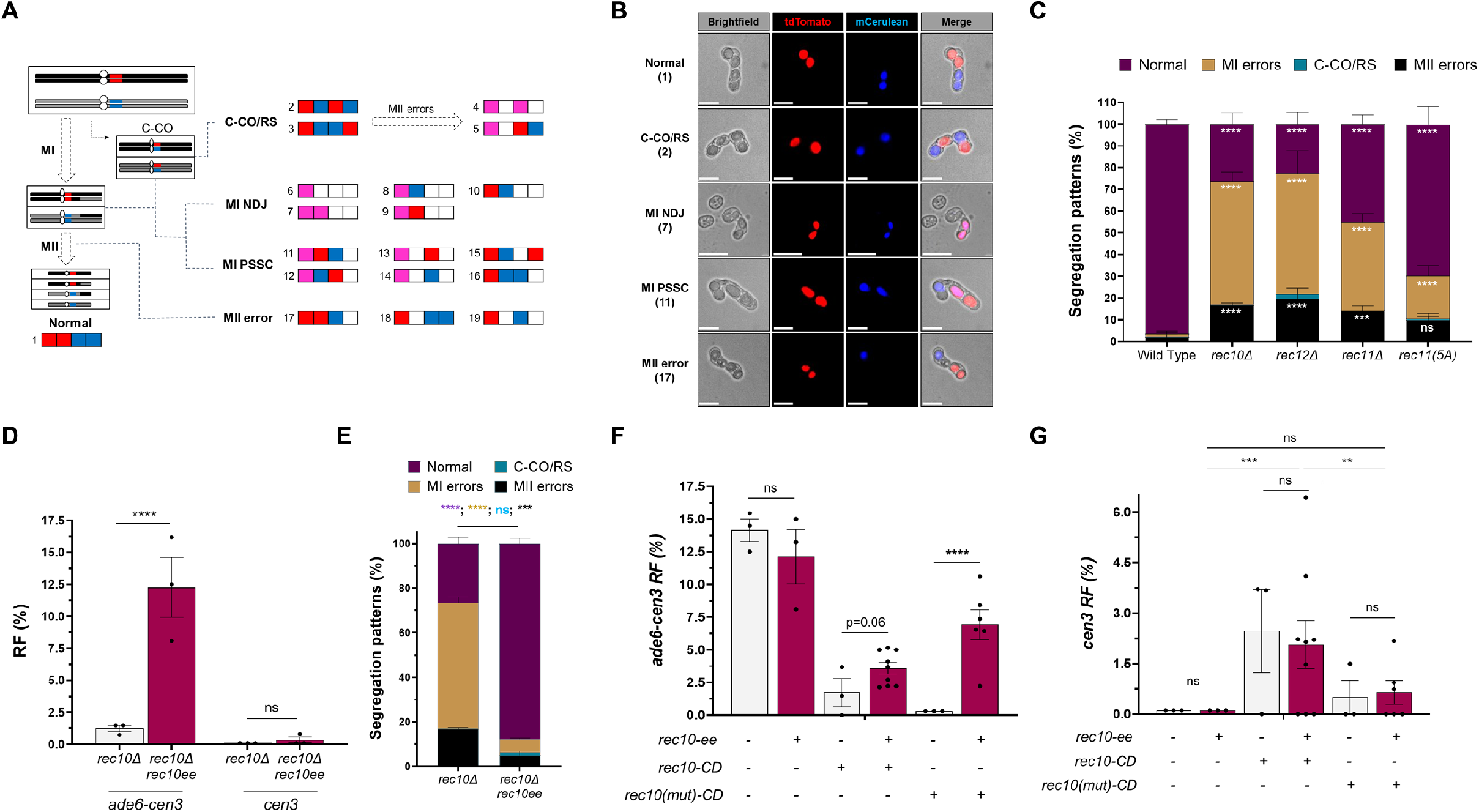
Lowered levels of chromosomal arm crossovers lead to meiosis I defects. **A**. Schematic representation of the fluorescence-based tetrad assay for segregation fidelity and the various types of patterns that can be obtained corresponding to the errors in specific stages of meiotic division. Patterns arising from mirroring each or both halves of the asci are not shown. **B**. Representative images for normal, centromeric crossover (C-CO) or reverse segregation (RS), meiosis I non-disjunction (MI NDJ), meiosis I premature separation of sister chromatids (MI PSSC) and meiosis II (MII) error types. One example from each of these error sub-types are shown here. Scale bars represent 10 μm. **C**. Plot showing frequency of normal, C-CO/RS, MI and MII segregation errors in wild type, *rec10Δ, rec12Δ, rec11Δ* and phosphomutant *rec11(5A)* genotypes. All data are mean ± SEM (n = 3-6 experiments), assaying 834, 509, 333, 489 and 364 tetrads for wild type, *rec10Δ, rec12Δ, rec11Δ* and *rec11(5A)*, respectively. **D**. Recombinant frequency (RF) across *ade6-cen3* arm interval and *cen3* in *rec10Δ* and *rec10Δ rec10ee* (ectopically expressed) genotypes. Data are mean ± SEM (n = 3 experiments), assaying 408 spore colonies for *rec10Δ* and *rec10Δ rec10ee*, each. **E**. Plot showing frequency of normal, C-CO/RS, MI and MII segregation errors in *rec10Δ* and *rec10Δ rec10ee* (ectopically expressed) genotypes. All data are mean ± SEM (n = 3 experiments), assaying 509 and 477 tetrads for *rec10Δ* and *rec10Δ rec10ee*, respectively **F-G**. RF across *ade6-cen3* (**F**) arm interval and *cen3* (**G**) in *wild type, rec10ee, rec10-CD, rec10-CD rec10ee, rec10(mut)-CD* and *rec10(mut)-CD rec10ee* genotypes. Data are mean ± SEM (n = 3-9 experiments), assaying 410, 412, 408, 1122, 401 and 631 spore colonies for *wild type, rec10ee, rec10-CD, rec10-CD rec10ee, rec10(mut)-CD* and *rec10(mut)-CD rec10ee*, respectively. ^****^p <0.0001; ^***^p <0.001; ^**^p <0.01; ^*^p <0.05; ns p >0.05. Data in panels C and E were analysed by Two-way ANOVA test, whereas Two-tailed Fisher’s exact test was used for data in panels D, F and G. The data for individual experiments are provided in Supplementary Tables 4, 5 and 6.

In meiosis I, non-disjunction (NDJ) of homologs, occurs when the homologs fail to separate towards opposite poles in MI or randomly segregate (Fig. 1A, tetrads 6-9, 1B and Suppl. Fig. 1A). Premature separation of sister chromatids (PSSC), implies sister chromatids separating in MI itself, towards opposite poles (Fig. 1A, tetrads 11-16, 1B and Suppl. Fig. 1A). MI PSSC could be likely generated due to biorientation of the sister kinetochores, facilitated by loss of mono-orientation factors such as Moa1 or loss of pericentric cohesion via Rec8-Psc3 complex. In meiosis II, errors during segregation could arise due to MII NDJ, wherein sister chromatids fail to separate or randomly segregate in the absence of the residual centromeric cohesion (Fig. 1A, tetrads 17-19; 1B and Suppl. Fig. 1A).

Another possible type of aberrant segregation can occur via reverse segregation (RS), where MI is equational (sister chromatids segregate) and meiosis II is reductional (homologs separate) owing to mono-orientation defects. This is different from a combination of two individual PSSC events as the frequency of occurrence of equational division due to reverse segregation is nearly 100 times higher than the expected frequency of a combination of two PSSC events as observed in human oocytes^24^. Alternatively, in an event of a crossover between the centromere and the fluorophore, the segregation profile will look identical to the reverse segregation profile (alternate colour pattern in each half of the ascus), upon normal segregation thereafter in MI and MII (Fig. 1A, tetrads 2,3; 1B and Suppl. Fig. 1A). In order to distinguish between the two, we additionally looked at the centromeric recombination frequency in such genotypes via genetic recombination assays. Since this assay cannot distinguish between RS and C-CO associated phenotypes by itself, we have named this category as an RS/C-CO phenotype. However, as will be discussed below, mutants disrupting mono-orientation fidelity that could promote reverse segregation, show distribution and type of errors very different from those that undergo C-COs. Hence, the consequences of C-COs and mono-orientation loss are easily distinguishable. Additionally, tetrads that undergo a C-CO followed by MI mis-segregation, would still show the same phenotype as normal MI NDJ or PSSC events. In contrast, if a C-CO is followed by normal MI segregation but erroneous MII segregation, a distinct profile will be observed that cannot result from any other source (Fig. 1A, tetrads 4,5; 1B and Suppl. Fig. 1A).

We also designed a system wherein addition of a third fluorophore at the other end of the centromere on one of the homologs might help in measurement of C-CO directly. But it increased the possible combinations of tetrads obtained, many of which, may still not be able to exhibit whether the mis-segregating chromosome had also undergone a C-CO (Suppl. Discussion).

### Absence of chromosomal arm crossovers exhibit meiosis I defects

In *S. pombe*, active exclusion of a functional Rec8-Rec11 cohesin complex at the pericentric heterochromatin prevents the pro-recombinogenic linear element (LinE) proteins such as Rec10 to be recruited at the centromeres^11,25^. We previously showed that forced recruitment of Rec10 to the pericentric heterochromatin by fusing it with the H3K9Me-binding chromodomain (CD) dramatically increases meiotic recombination frequency across fission yeast centromeres *cen1* and *cen3*^11^. However, this concomitantly led to a significant decrease in recombination frequencies across chromosomal arm intervals, due to significant redistribution of Rec10 to the centromeric and other heterochromatin regions from the chromosomal arms.

Since arm crossovers have been known to aid in proper chromosomal segregation, their absence would elevate defects during meiotic divisions^20^. In order to confirm and determine the subtype of errors upon reduced arm crossovers, we performed our assay in mutants’ defective at different steps of DSB initiation. Phosphorylation of the meiotic cohesin subunit Rec11, helps recruit Rec10, which further activates the Rec12 complex to initiate DSB formation in *S. pombe*^26^. We used *rec10*Δ, *rec12*Δ, *rec11*Δ and a phosphodeficient mutant *rec11(5A)*, to study the segregation defects using the tetrad assay and evaluated asci with four spores. The error profiles showed ∼40-60% MI errors, especially in *rec10*Δ, *rec12*Δ *rec11*Δ strains, followed by ∼15-20% MII errors (Fig. 1C and Suppl. Fig. 1B). *rec11(5A)* showed overall lesser MI errors as compared to *rec11*Δ possibly due to intact arm cohesion-associated functions (Fig. 1C and Suppl. Fig. 1B). In addition, the *rec11(5A)* mutant still retained significant recombination frequency at the *ade6-arg1* interval compared to *rec11*Δ that had completely abolished recombination, suggesting possibly additional phosphorylatable sites on Rec11 apart from the 5 modified sites (Suppl. Fig. 3B). Within the MI population, all the genotypes showed predominantly higher NDJ events as compared to PSSC events (Fig. 1C and Suppl. Fig. 1B). The increased MII errors in these strains may be due to a previously reported non-topoisomerase catalytic activity of Rec12, which is required for proper segregation fidelity of sister-chromatids in meiosis II^27^. Hence, in the absence of Rec12 or its recruitment and activation by Rec11-Rec10 complex, errors in MII may also happen.

In order to study the specific effects of C-COs generated by Rec10-CD, we wanted to establish a genetic background where C-COs may be the only source of segregation errors and restore arm crossovers to abolish the consequent segregation defects. Hence, we first checked whether over-expressing Rec10 ectopically in a *rec10*Δ genetic background could rescue the recombination and segregation defects. The presence of ectopic copy of Rec10 via plasmid and overexpression of Rec10 was confirmed via PCR and immunoblotting (Suppl. Figs. 1C and 1D). The recombination frequency (RF) across the chromosomal arm interval *ade6-cen3* in *rec10*Δ was 1.2%, whereas in the presence of *rec10ee*, the RF increased to 12.25%, which is similar to wild type indicating complete rescue upon ectopic expression of Rec10 (Figs. 1D, 1F). The *cen3* recombination stayed repressed in both the genetic backgrounds (Fig. 1D). For segregation, the error profiles in the *rec10*Δ strains prominently showed MI errors, wherein NDJ events were 39.6% while the PSSC events were 16.8% (Fig. 1E). However, upon ectopic expression of Rec10 in the *rec10*Δ background, the total segregation errors came down by ∼90%, indicating that the ectopically expressed Rec10 was able to rescue the reduced arm recombination-associated segregation defects in *rec10*Δ background (Fig. 1E).

Next, we confirmed the presence of ectopic copy of Rec10 in wt, *rec10-CD* and *rec10(mut)-CD* genotypes and measured arm and centromere recombination in these genotypes (Suppl. Fig. 1D and Figs. 1F, 1G). No significant changes were observed in the RF at the *ade6-cen3* interval in wild type expressing ectopic Rec10 as expected (Fig. 1F). However, there was an increase in the arm RF in both the *rec10-CD* (3.7% compared to 1.7%) and *rec10(mut)-CD* (6.5% compared to <0.2%) strains in the presence of *rec10ee* suggesting rescue in recombination, although not complete (Fig. 1F). Since, C-COs are associated with increased chromosomal mis-segregation and aneuploidy, they may result in a higher degree of spore inviability. This could explain why *rec10(mut)-CD* (which does not have high C-COs) shows a better rescue of arm RF upon Rec10 overexpression. Importantly, there was a significant increase in recombination across *cen3* in the presence of *rec10-CD* (2.5%) as expected, which didn’t change upon ectopic expression of *rec10ee* (Fig. 1G). This clearly indicated that introduction of ectopic copy of *rec10*, could rescue arm RF, without affecting *cen* RF and hence could be used to study the effects of C-COs on meiotic chromosome segregation.

### Increased centromere-proximal crossovers reduce spore viability

Since, ectopic expression of Rec10 did not completely rescue arm recombination, in the C-CO proficient strain *rec10-CD*, unlike in *rec10*Δ background, we wondered whether presence of C-COs could be leading to spore death due to increased chromosomal mis-segregation and aneuploidy that may result in an underestimation of recombination frequencies. In order to check this, we stained the asci with Propidium Iodide (PI), which is known to stain dead cells that have lost their membrane integrity but is efficiently excluded from healthy cells. This method has been used previously, to estimate spore death in meiotic asci as well^28,29^. Meiotic asci in *rec10*Δ showed a high degree of PI-stained aberrant spores, indicating poor spore health, which got rescued upon ectopic expression of Rec10 (Suppl. Fig. 2A and Figs. 2A, 2B). We observed aberrant asci with 1, 2, 3 or all 4-spores PI stained, exhibiting the spectrum of potentially inviable spores in absence of Rec10 (Fig. 2A, 2B). Similarly, we observed rescue in the number of asci with aberrant PI-stained spores for both *rec10-CD* and *rec10(mut)-CD* upon ectopic Rec10 expression (Suppl. Fig. 2A). The percentage of normal and aberrant asci for *rec10-CD* after rescue with *rec10ee* was similar to that of *rec10*Δ, suggesting that the arm recombination-mediated viability defects were being completely rescued (Fig. 2B and Suppl. Fig. 2A). However, despite this similarity, the incomplete rescue of arm recombination seen above, may be attributed to the additional defects conferred by the presence of C-COs that may further affect spore viability leading to underestimation of recombinants in the genetic assay. The rescue of spore viability in *rec10(mut)-CD* wasn’t at par with that of the *rec10*Δ, which could explain why the arm recombination frequency didn’t quite reach wild type levels (Figs. 1D and 2B).

**Figure 2.**
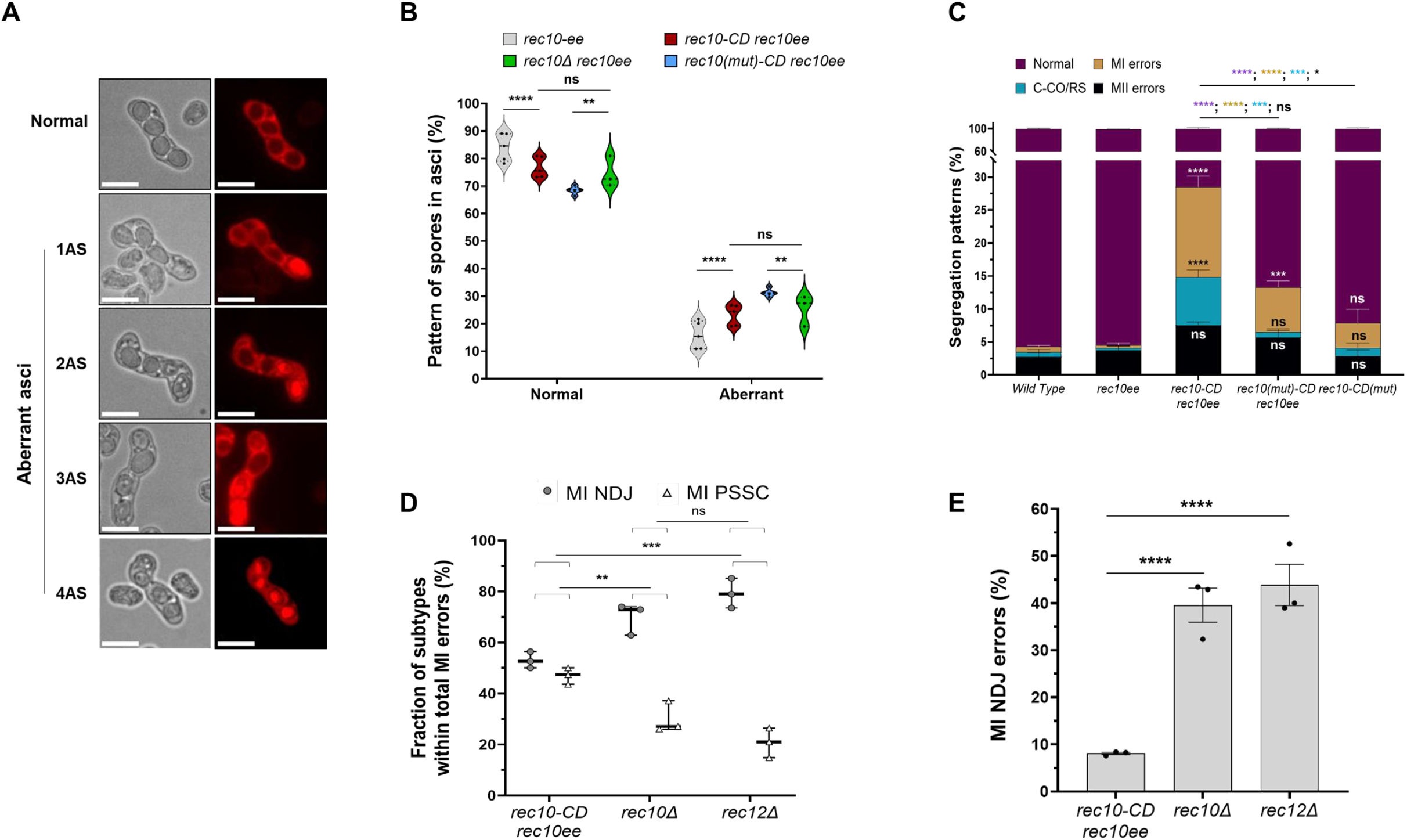
Elevated pericentric crossovers lead to increased meiosis I segregation defects and spore inviability. **A**. Representative images for normal and aberrant asci after staining with propidium iodide (PI). The aberrant asci contain 1 – 4 PI-stained “dead” spores. Scale bars represent 10 μm. **B**. Graph showing distribution of normal and aberrant asci (containing 1-4 PI-stained spores) in *rec10ee, rec10-CD rec10ee, rec10(mut)-CD rec10ee* and *rec10Δ rec10ee* genotypes. All data are mean ± SEM (n = 3-5 experiments), assaying 1059, 1580, 818 and 882 tetrads for *rec10ee, rec10-CD rec10ee, rec10(mut)-CD rec10ee* and *rec10Δ rec10ee*, respectively. **C**. Plot showing frequency of normal, centromeric crossover (C-CO) or reverse segregation (RS), meiosis I (MI) and meiosis II (MII) segregation errors in wild type, *rec10ee, rec10-CD rec10ee, rec10(mut)-CD rec10ee* and *rec10-CD(mut)* genotypes. The significance values within the bars in the graph are with respect to *rec10ee*. All data are mean ± SEM (n = 3 experiments), assaying 890, 367, 479, 517 and 560 tetrads for wild type, *rec10ee, rec10-CD rec10ee, rec10(mut)-CD rec10ee* and *rec10-CD(mut)*, respectively. **D**. Graph showing distribution of MI non-disjunction (NDJ) and MI premature separation of sister chromatids (PSSC) events among the total MI errors in *rec10-CD rec10ee, rec10Δ* and *rec12Δ* genotypes. **E**. Bar graph showing absolute percentage of MI NDJ error events in *rec10-CD rec10ee, rec10Δ* and *rec12Δ* genotypes. ^****^p <0.0001; ^***^p <0.001; ^**^p <0.01; ^*^p <0.05; ns p >0.05. Data in panels B-D were analysed by Two-way ANOVA test, whereas Two-tailed Fisher’s exact test was used for data in panel E. The data for individual experiments are provided in Supplementary Tables 4, 5 and 7.

### Increased centromere-proximal crossovers are associated with elevated errors at meiosis I

In order to study the deleterious effects of increased C-COs on meiotic chromosome segregation, the assay was performed in the *rec10-CD rec10ee* genotype with confirmed ectopic expression (ee) of Rec10 (Suppl. Fig. 1E). *Rec10-CD* alone showed only 41.1% tetrads with normal segregation, whereas most of the errors were contributed by abnormal MI segregation (33.3%), followed by 17.7% MII errors (Suppl. Fig. 2B). We also observed a significant fraction of tetrads (8.1%) showing C-COs phenotype confirming the activation of meiotic recombination events at the otherwise repressed centromeres (Fig. 2C and Suppl. Fig. 2B). When Rec10 was ectopically overexpressed in the presence of *rec10-CD*, the percentage of normal segregation increased to 72.2%, without significantly affecting frequency of C-COs (7.3%) (Suppl. Fig. 2B and Fig. 2C). Importantly, the MI errors were still significantly high in the presence of *rec10-CD rec10ee* (13.8%), especially when compared to a *rec10(mut)-CD rec10ee* (6.8%), where the C-COs were nearly abolished and MII errors largely remained the same (Fig. 2C).

Ectopic Rec10 expression was able to dramatically reduce all the error classes even in presence of *rec10(mut)-CD*, suggesting that the lowered arm recombination-associated defects were rescued similar to *rec10*Δ (Suppl. Fig. 2B). We also observed that most of the errors observed in presence of *rec10-CD* were lost upon using a *rec10-CD(mut)* allele harboring a W104A mutation in the CD domain, making it unable to bind to H3K9 methylated heterochromatin^30^ (Fig. 2C). Since the errors are largely rescued in case of pericentric targeting of Rec10 mutant (*rec10(mut)-CD rec10ee*) or a CD mutant [*rec10-CD(mut)*], the errors observed in *rec10-CD rec10ee* are plausibly linked to C-CO activation by pericentric recruitment of functional Rec10.

Hence our results suggest that elevated frequency of C-COs is strongly associated with increased meiosis I errors. Interestingly, within this class, we observed a slight bias towards MI NDJ as compared to MI PSSC events (Fig. 2D and Suppl. Fig. 2C). Out of the total tetrads showing MI errors, 54.4% were contributed by MI NDJ in *rec10-CD rec10ee*, while MI PSSC occurred in 45.6% tetrads (Fig. 2D). Overall, these data show that C-COs in *S. pombe* disrupt accurate disjunction of homologous chromosomes during meiosis I. Interestingly, both absence of COs in the arms and presence of COs at the centromeres show defects in meiosis I, which appear counterintuitive but support the necessity for preserving centromeric recombination repression across species. However, mechanistically MI NDJ events in both these contexts should be different. Since both *rec10*Δ and *rec12*Δ are completely recombination deficient, the non-disjunction of homologs in meiosis I may not reflect the classical inability of the chromosomes to separate and segregate into the same nucleus. Loss of physical connections between homologs would result in random segregation of the chromosomes with both the sister chromatids still together, unlike PSSC. This is also supported by the nearly 50% MI NDJ errors seen in *rec10*Δ and *rec12*Δ that reflect random homolog segregation and much reduced PSSC events within the total MI errors, unlike in the presence of C-COs (Figs. 2D, 2E). Therefore, these may be better classified as MI “NDJ-like” events in such recombination-deficient mutants.

### Loss of centromeric cohesion does not lead to increase in meiosis I errors

Since a significant fraction of the MI errors are attributed to premature separation of sister chromatids, we wondered if the elevated MI errors in case of increased C-COs could be due to loss of pericentric cohesion triggered during the process of Holliday Junction formation or resolution of recombination intermediates. If so, the errors generated in genotypes that are unable to establish, maintain or preserve pericentric cohesion, should mimic those generated during C-COs formation (Fig. 3A). The pericentric cohesin complex is different from that of the arms in having Psc3 as the HAWK subunit as compared to Rec11, while the trimeric ring Psm1-Psm3-Rec8 stays the same^25^. This complex is also essential for maintaining chromosomal segregation fidelity in MI, since the centromeric cohesion needs to be preserved at anaphase I to keep sister chromatids attached to each other and favor disjunction of homologs. Weakened centromeric cohesion, which is elevated in aged mammalian oocytes, increases rates of meiotic aneuploidy, predominantly PSSC events ^31^.

**Figure 3.**
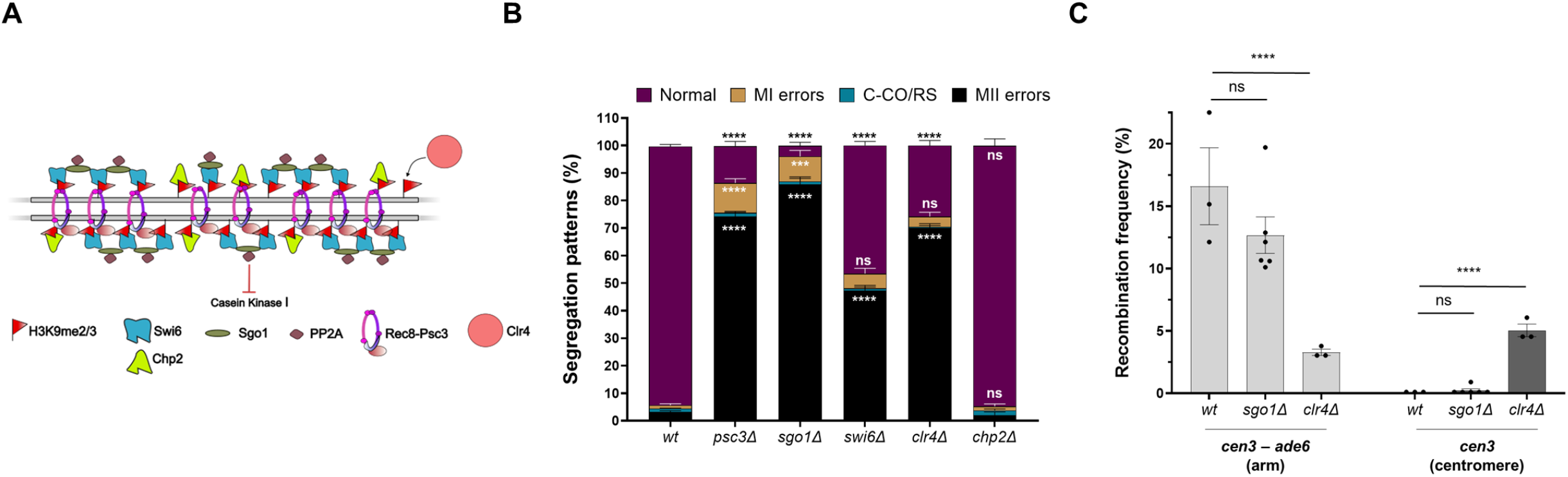
Loss of centromeric cohesion and heterochromatin result in meiosis II errors. **A**. Schematic representation of the different pericentromere localising proteins, mutants of which were used to disrupt the centromeric cohesion and the pericentric heterochromatin. **B**. Plot showing frequency of normal, centromeric crossover (C-CO) or reverse segregation (RS), meiosis I (MI) and meiosis II (MII) segregation errors in wild type, *psc3Δ, swi6Δ, sgo1Δ, clr4Δ* and *chp2Δ* genotypes. Data are mean ± SEM (n = 3 experiments), assaying 832, 364, 377, 415, 215 and 615 tetrads for wild type, *psc3Δ, swi6Δ, sgo1Δ, clr4Δ* and *chp2Δ*, respectively. **C**. Recombinant frequency (RF) across *cen3-ade6* arm interval and *cen3* in wild type, *sgo1Δ* and *clr4Δ* genotypes. Data are mean ± SEM (n = 3-6 experiments), assaying 374, 683 and 462 spore colonies for wild type, *sgo1Δ* and *clr4Δ*, respectively. ^****^p <0.0001; ^***^p <0.001; ns p >0.05. Data in panel B were analysed by Two-way ANOVA test, whereas Two-tailed Fisher’s exact test was used for data in panel C. The data for individual experiments are provided in Supplementary Tables 9 and 10.

First, we tested meiotic segregation fidelity in *psc3*Δ cells, which can survive if the meiotic cohesins Rec8 and Rec11 are ectopically expressed in vegetative cells^25^. Upon completion of meiosis in the absence of Psc3, a dramatic 74.2% tetrads showed MII errors due to random sister-chromatid separation arising from the absence of the critical centromeric cohesion (Fig. 3B). There were only 13.4% normal tetrads and merely 10.8% tetrads showing MI errors. As expected, there were no significant C-COs seen, supporting genetic recombination data determined previously^11^ (Fig. 3B). This overwhelming skew toward MII errors points towards the roles of Psc3, as part of the centromeric cohesin complex, mainly during segregation of sister-chromatids. The protection of this centromeric cohesin population is facilitated by the Sgo1-PP2A complex that binds to and prevents cleavage of the Rec8-Psc3 complex by inhibiting its phosphorylation by the CK1 kinase^32^ (Fig. 3A). As expected, in *sgo1*Δ, 85.8% tetrads showed MII errors, with close to no normal segregation, which interestingly is a much more severe phenotype than *psc3*Δ, reiterating that failure to preserve the centromeric cohesion caused mainly meiosis II errors (Fig. 3B). *sgo1*Δ has normal chromosomal arm recombination frequency across the measured *cen3-ade6* genetic interval, compared to wild type and no increase in recombination across *cen3*, confirming that the segregation defects were not due to aberrant recombination (Fig. 3C). Deletion of the catalytic subunit of the PP2A complex, *ppa2*Δ did not phenocopy *sgo1*Δ and was mostly indistinguishable from wild type, except for a relatively lower but significant increase in MII errors (11.6%) as compared to *psc3*Δ and *sgo1*Δ (Suppl. Fig. 2D). This is in agreement with a previous report that deletion of the Par1-B subunit of the PP2A complex, and not the catalytic subunit Ppa2, showed segregation errors comparable to *sgo1*Δ^33^. This may also suggest contribution of other phosphatases apart from PP2A that could preserve Rec8 from cleavage in its absence. Overall, disruptions in centromeric cohesion dramatically elevated the errors in sister-chromatid segregation in MII, much more than affecting homolog disjunction in MI. In all these cases, since the chromosomal arm cohesion by Rec8-Rec11 complex is retained and the crossover formation in the arm is unhindered, it can be expected that the homolog disjunction during MI stays unaffected.

### Disruption of pericentric heterochromatin elevates meiosis II errors unlike centromere-proximal crossovers

Since, Rec10-CD activates C-COs in the pericentric heterochromatin, we checked if perturbed heterochromatin function is the contributing factor towards the increased C-CO associated MI errors. Heterochromatin protein HP1 orthologs, Swi6 and Chp2, recognize and bind the histone H3K9me epigenetic marks and help in heterochromatin assembly and silencing functions (Fig. 3A). Additionally, Swi6 promotes deposition of the Rec8-Psc3 cohesin complex at the pericentric heterochromatin to facilitate sister-chromatid cohesion^34^. On the other hand, Chp2 promotes histone deacetylation during heterochromatin assembly by recruiting the SHREC complex^35^. We tested whether disruption of heterochromatin activity by removing either Swi6 or Chp2, affected segregation fidelity and observed an increased frequency of MII errors upon *swi6*Δ (47.2%), which was much less severe than both *psc3*Δ and *sgo1*Δ (Fig. 3B). Both C-CO and MI errors were not different among these three genotypes. Interestingly, in *chp2*Δ the effects were unlike its paralog Swi6 and nearly wild type, suggesting that disruption of heterochromatin gene silencing functions via Chp2, does not play any role in maintaining chromosomal segregation fidelity (Fig. 3B). In contrast, affecting heterochromatin’s role in centromeric cohesin deposition via Swi6, severely compromises segregation fidelity, thereby providing evidence for another non-overlapping function of the HP1 paralogs Swi6 and Chp2.

Clr4 is the only known histone methyl transferase in *S. pombe* and hence its deletion results in absence of H3K9 methylation, and consequently leads to abolishment of heterochromatin. *clr4*Δ also results in low levels of Swi6 and consequently low pericentric cohesins^34^. Since *clr4*Δ can also lead to derepression of C-COs^36^, we confirmed the increase in centromeric recombination events, and also notably, a concomitant decrease in arm recombination across the *cen3-ade6* interval (Fig. 3C). This was similar to what we observed in the C-CO proficient *rec10-CD rec10ee* cells (Figs. 1F, 1G). In contrast, the segregation profiles showed a significant increase in MII errors in *clr4*Δ, as opposed to the elevated MI errors observed in the *rec10CD rec10ee* background (Fig. 3B). The increase in MII errors is similar to that observed in *swi6*Δ and *psc3*Δ, due to the lowered pericentric cohesion. Hence, we can infer that loss of functional heterochromatin cannot explain the elevated MI errors due to C-COs, since such mutants also show predominantly MII errors, which is in agreement with the previously described low levels of Rec8-Psc3 cohesins and consequently centromeric cohesion in *swi6*Δ and *clr4*Δ genotypes^34^.

Overall, it appears that the deleterious segregation effects due to loss of pericentric cohesins can outweigh the errors due to abnormal recombination (increased C-COs or absence of arm COs) and switch the mis-segregation types to predominantly MII errors. This argues that if C-COs would be considerably affecting the pericentric cohesion or heterochromatin establishment, we would have observed a high MII error phenotype, which is not observed. Hence, we infer that C-CO events do not significantly disrupt pericentric cohesion directly.

### Residual pericentric cohesins at MI contribute towards homolog non-disjunction by centromeric-crossovers

A major fraction of the MI errors due to C-COs by *rec10-CD rec10ee* were due to non-disjunction events, which appear to be mechanistically different from those seen in *rec10*Δ or *rec12*Δ mutants that are devoid of arm COs and maybe distributing homologs randomly in MI daughter nuclei (Figs. 2D, 2E). One of the major differences between the arm and centromeres is the cohesin dynamics during late prophase I. The centromeric cohesin population is protected till sister-chromatid separation at MII, while the arm cohesins are proteolytically cleaved at end of MI^37^. Removal of cohesins distal to the chiasma is necessary to facilitate separation of the homologs that underwent recombination^4,38^. Could the persisting centromeric cohesins promote the observed MI NDJ events, when C-COs occur that are otherwise repressed?

In the absence of the centromere-specific cohesin subunit Psc3, there is a high frequency of MII errors and no activation of C-COs, as established above (Fig. 3B). In this background, we planned to activate C-COs by targeting the chromosome arm specific and recombinogenic Rec8-Rec11 complex via chromodomain fusions, which previously has been shown to derepress pericentric recombination^11^ (Suppl. Fig. 3A). We compared this with a *rec11(5A)-CD* mutant that is recombination-deficient at the chromosomal arms. An ectopic copy of the wild type Rec11 (*rec11ee*) was expressed in these genotypes to maintain a robust frequency of chromosomal arm crossovers and rule out any MI segregation defects due to reduced arm recombination (Suppl. Fig. 3B). We confirmed the rescue of MI defects arising in presence of *rec11*Δ or *rec11(5A)*, as seen above, via the ectopic expression of the *rec11* allele (Suppl. Fig. 3C). Interestingly, neither targeting Rec8-Rec11 complex to pericentric regions via CD nor presence of *rec11ee* were able to rescue the severe MII defects in *psc3*Δ (Suppl. Fig. 3D). However, we also didn’t observe any significant differences in the C-CO or MI errors in *psc3*Δ *rec8-CD rec11-CD rec11ee* compared to *psc3*Δ *rec8ee rec11ee* (Suppl. Fig. 3D). We suspect that due to the high levels of MII errors and consequences of loss of centromeric Psc3 on sister-chromatid cohesion, we might be unable to detect the majority of the C-CO events and consequent MI errors. Moreover, the phosphodeficient mutant *rec11(5A)-CD* in this genotypic background, didn’t reduce the 8-11% of MI errors seen in the *psc3*Δ *rec8-CD rec11-CD rec11ee* strain, probably due to the presence of an ectopic copy of wild type Rec11 that may also access the centromere along with Rec8-CD (Suppl. Fig. 3D). One striking observation of the MI errors in the absence of Psc3, was a conspicuous change in the distribution of the fraction of MI error subtypes (Fig. 4A and Suppl. Fig. 3E). There was a dramatic increase in MI PSSC events as compared to MI NDJ events in *psc3*Δ genotypes, which was in stark contrast to that observed in *psc3+ rec10-cd* strains as shown above (Compare Fig. 4A and Fig. 2D).

**Figure 4.**
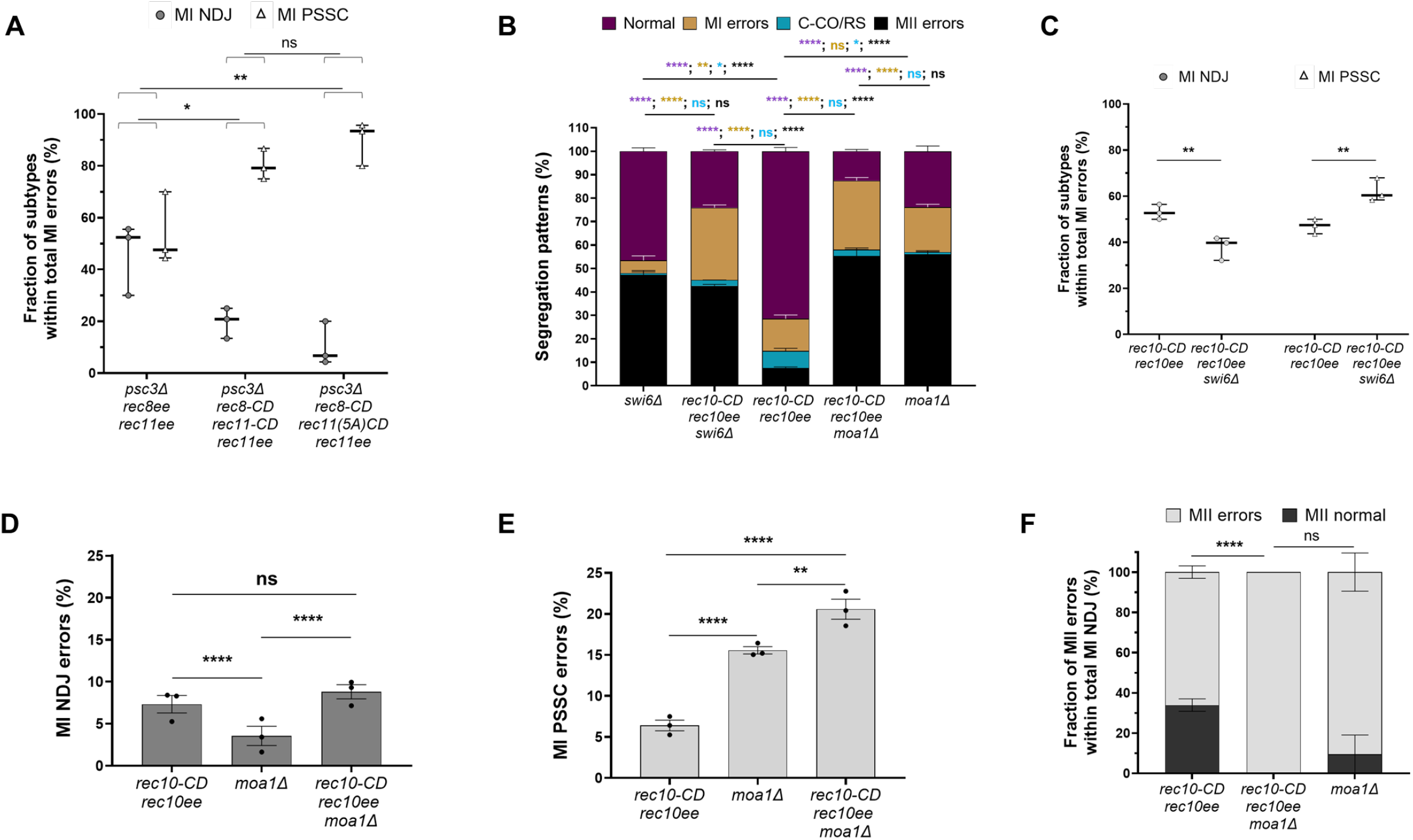
Reduced centromeric cohesion and mono-orientation fidelity alters the subtype of meiosis I errors in the presence of increased centromeric crossovers. **A**. Graph showing distribution of MI non-disjunction (NDJ) and MI premature separation of sister chromatids (PSSC) events among the total MI errors in *psc3Δ rec8ee rec11ee* (ectopically expressed), *psc3Δ rec8-CD rec11-CD rec11ee* and *psc3Δ rec8-CD rec11(5A)-CD rec11ee* genotypes. All data are mean ± SEM (n = 3 experiments), assaying 364, 477 and 647 tetrads for *psc3Δ rec8ee rec11ee, psc3Δ rec8-CD rec11-CD rec11ee* and *psc3Δ rec8-CD rec11(5A)-CD rec11ee*, respectively. **B**. Plot showing frequency of normal, centromeric crossover (C-CO) or reverse segregation (RS), meiosis I (MI) and meiosis II (MII) segregation errors in *swi6Δ*, r*ec10-CD rec10ee swi6Δ, rec10-CD rec10ee*, r*ec10-CD rec10ee moa1Δ* and *moa1Δ* genotypes. **C**. Graph showing distribution of MI NDJ and MI PSSC events among the total MI errors in r*ec10-CD rec10ee* and *rec10-CD rec10ee swi6Δ* genotypes. **D**. Bar graph showing absolute percentage of MI NDJ error events in *rec10-CD rec10ee, moa1Δ* and r*ec10-CD rec10ee moa1Δ* genotypes. **E**. Bar graph showing absolute percentage of MI PSSC events in *rec10-CD rec10ee, moa1Δ* and r*ec10-CD rec10ee moa1Δ* genotypes. **F**. Graph showing fraction of MI NDJ events with normal and erroneous MII segregation in *rec10-CD rec10ee*, r*ec10-CD rec10ee moa1Δ* and *moa1Δ* genotypes. All data are mean ± SEM (n = 3 experiments), assaying 479, 713, 377, 635 and 843 tetrads for *rec10-CD rec10ee*, r*ec10-CD rec10ee swi6Δ, swi6Δ*, r*ec10-CD rec10ee moa1Δ* and *moa1Δ*, respectively. ^****^p <0.0001; ^**^p <0.01; ^*^p <0.05; ns p >0.05. Data in panel B were analysed by Two-way ANOVA test, whereas Two-tailed Fisher’s exact test was used for data in panels A, C-F. The data for individual experiments are provided in Supplementary Tables 5 and 9.

We speculated that lowering the pericentric cohesin population might aid in reducing the MI NDJ events occurring due to unresolved centromeric crossovers between homologs. In order to further strengthen this observation, we employed *swi6*Δ, as a way to reduce the cohesins in the vicinity of the C-COs generated by *rec10-CD rec10ee*, since overall, the frequency of MI and MII errors was lesser in *swi6*Δ compared to *psc3*Δ^34^ (Fig. 3B). As expected, we observed a significant increase in the total MI errors due to *rec10-CD rec10ee* in *swi6*Δ as compared to *swi6*Δ and *rec10-CD rec10ee* alone, which could also be due to more recruitment of Rec10-CD to the H3K9me heterochromatin, in absence of the competing Swi6 (Fig. 4B). Between *rec10-CD rec10ee* ± *swi6*Δ, MI PSSC events increased significantly and not MI NDJs (Fig. 4B and Suppl. Fig. 4A). However, if we compare the individual MI NDJ and MI PSSC fractions among the total MI errors, we continued to observe a significantly marked reduction in the MI NDJ events and a concomitant increase in levels of PSSC phenotype (Fig. 4C and Suppl. Fig. 4B). This suggested that presence of the protected population of pericentric cohesins during anaphase I, may hinder the timely resolution of the C-COs, resulting in the nondisjunction of the homologs, which gets lowered upon reducing cohesion (Suppl. Fig. 4B). Additionally, our data indicates that C-COs may also interfere with the appropriate monopolar attachment of the sister chromatids. Loss of mono-orientation can result in the tendency of atleast one pair of sister chromatids to split at MI that gets exacerbated upon reducing pericentric sister chromatid cohesion (Suppl. Fig. 4B).

### Centromeric crossovers may additionally disrupt mono-orientation of homologs at MI

Moa1 is integral for mono-orientation of homologs in MI and its loss results in biorientation of sister chromatids due to loss of active core centromeric cohesion^39^. Interestingly, *moa1*Δ shows maximal MII errors (56%) similar to *swi6*Δ and 19.1% total MI errors (Fig. 4B). The near absence of C-CO/RS with normal MII phenotype (Fig. 1A, tetrads 2 and 3) in *moa1*Δ, strongly argues that the corresponding error pattern observed in *rec10-cd rec10ee* arises from C-COs with intact pericentric cohesion, since equational segregation in MI would require removal of sister-chromatid cohesion to ultimately allow the separation of chromosomes. Among the total MI errors, *moa1*Δ showed a higher percentage of MI PSSC events compared to NDJ, suggesting biorientation and separation of atleast one pair of sister chromatids at MI itself (Fig. 4D, 4E). Upon addition of C-COs via Rec10-CD, there was no further increase in the frequency of MI NDJ events, compared to *rec10-CD rec10ee* alone, suggesting that the co-migration of the homologs in the presence of C-COs couldn’t be further alleviated by loss of mono-orientation (Fig. 4D). It is plausible that the unresolved centromere-proximal crossovers result in pulling the two homologs into the same nucleus, despite loss of mono-orientation fidelity, leading to NDJ events. Conversely, the PSSC events showed an additive effect in the double mutant *rec10-CD rec10ee moa1*Δ when compared to either mutant alone (Fig. 4E). This indicates that C-COs may independently disrupt accurate mono-orientation of the homologs leading to premature separation of sister chromatids atleast in half of the homologs, resulting in MI errors. Interestingly, when we look within the MI NDJ events, they can be sub-classified into those tetrads with normal MII (Figs. 1A, tetrad 7, 1B), MII error involving one homolog (Fig. 1A, tetrads 8 and 9; Suppl. Fig. 1A) and complete MII errors in both homologs (Fig. 1A, tetrads 6 and 10; Suppl. Fig. 1A). We find that the MI NDJ errors in C-COs alone have a significantly higher frequency of normal MII as compared to C-COs *moa1*Δ or *moa1*Δ alone (Fig. 4F). This is also true for frequency of MII NDJ in one homolog being significantly higher than MII NDJ in both homologs in presence of C-COs alone, as compared to when *moa1*Δ is present (Suppl. Fig. 4C). This argues for the idea that C-COs may actually end up protecting the bivalent from age-related consequences such as loss of cohesion and kinetochore- orientation defects from PSSC events. Such stabilized but compromised tetrads in meiosis I would then exhibit NDJ events at MII among the older gametes, as has been hypothesized previously^15^.

## Discussion

Centromere-proximal crossovers are known to be repressed across a wide range of species and by multiple mechanisms. This repression is highly correlated with increased chromosomal mis-segregation and gametic aneuploidy, which reflects the reason for its highly conserved nature. Despite this, our current understanding of how C-COs cause mis-segregation lacks depth. Models to explain the increased MII NDJ upon increased centromere-proximal exchange, previously reported in mammalian oocytes, involve entangled chromosomes at MI leading to the bivalent passing intact into the MII nucleus and dividing reductionally giving rise to sister-chromatids in the same gametes^14,20^. Hence, the C-CO-mediated defect occurring at MI gets visualized as an MII error. However, the origin of the error should still be considered as MI. Alternatively, resolution of C-COs may lead to loss of pericentric cohesins leading to PSSC and the sister chromatids may or may not comigrate at MI, but consequently will have a 50% chance of randomly moving into the same nucleus during MII, resulting in an apparent MII NDJ event^14,15,20^. Hence, it appears that both of these models indicate that despite the source of the segregation errors being at MI, the phenotypic classification of the C-CO mediated errors has been historically made as MII non-disjunction, which can be misleading while elucidating the molecular basis of how C-COs cause chromosomal mis-segregation.

Our data in *S. pombe* exhibits that presence of C-COs predominantly leads to MI errors with a slight bias towards NDJ events over PSSC. Moreover, we demonstrate using various mutants that any perturbations to the pericentric cohesins directly results in MII errors, with intact MI segregation. This supports the model that the persisting population of pericentric cohesins can prevent the efficient resolution of chiasma, resulting in entangled homologs that fail to separate. Alternatively, homologs can randomly segregate and move into the same nucleus at MI due to loss of physical contact via crossovers. Such NDJ-like events were observed in recombination-deficient mutants such as *rec12*Δ and *rec10*Δ. Removal of cohesins distal to the site of crossovers is important for the resolution of the chiasma and the protected pericentric cohesion may prevent the resolution of crossovers, that lead to MI NDJ^40,41^. A previous report showing an MII oocyte having two homologous chromosomes connected at the centromeres with centromeric REC8 and SGO1 encompassing both sets of sister kinetochores, suggests that cytologically such a scenario where the homologs remain stuck to each other is possible that can contribute to gametic aneuploidy^40^.

We attempted to test the effect of removing centromeric cohesion while concomitantly elevating C-COs by targeting pro-recombinogenic Rec8-Rec11 cohesin complex to the pericentric regions via chromodomain fusions in *psc3*Δ^11^. Here, we noticed a high frequency of MII errors similar to *psc3*Δ, even though C-COs were elevated, due to the dominant effect of Psc3 removal. The striking observation, however, was the dramatic dip in the frequency of MI NDJ among the total MI errors and its redistribution into the MI PSSC category. This supports the idea that loss of centromeric cohesion may facilitate better chiasma resolution at the pericentric regions and aid in the separation of homologs. We further confirmed these observations in a *rec10-CD swi6*Δ background, where the levels of pericentric cohesion would be reduced, without any changes to the arm COs or distribution of the native cohesin population. We found that similar to the *psc3*Δ background, there was a significant redistribution of the subtypes within the total MI errors. We strongly suspect that homolog nondisjunction appears to be a driving factor for errors, since it might overshadow any potential loss in monopolar attachment that may occur in presence of C-COs, due to homologs getting entangled and moving towards the same pole. However, in a pericentric environment with lowered cohesin and thereby improved chiasma resolution, the biorientation of sister chromatids may be more pronounced, resulting in the marked increase in PSSC events.

Removal of mono-orientation protein Moa1 in presence of *rec10-CD rec10ee* did not alter the MI NDJ frequency, suggesting that even upon mono-orientation loss, C-COs still cause homolog nondisjunction, further proving its dominant effect. On the contrary, *moa1*Δ and *rec10-CD rec10ee* together show an additive increase in PSSC events, suggesting that each can cause biorientation of atleast one pair of sister-chromatids at meiosis I itself, independently. Mere loss of pericentric cohesion cannot explain PSSC events, as they primarily show defects in sister chromatid separation at MII. Additionally, this is supported by observations in a recent study, wherein presence of single chromatids was found to be rare in MI arrested human oocytes irrespective of the donor age, indicating that cohesion in chromosomal arms is enough to keep the sister chromatids together and centromeric cohesion becomes indispensable only after removal of chromosomal arm cohesins at anaphase I^42^.

Overall, our study provides sufficient evidence to support the following model for the long known, but poorly understood reason for how the centromere-proximal crossovers cause disruption of proper chromosome segregation, despite arm crossovers being critical for the same (Fig. 5). C-COs due to their position at the centromeres are unable to get resolved in the absence of cohesin removal during anaphase I, resulting in entanglement of the recombining homologs along with the paired sister-chromatids and thereby segregating into the same nucleus upon completion of MI. This gives rise to the classical “nondisjunction” phenotype of the homologs (Fig. 5). Additionally, we suspect that C-COs may also interfere with the proper monopolar attachment of either one or both pairs of sister chromatids. When it occurs for one pair of sister chromatids, it results in a PSSC event. However, if it occurs on both it would lead to reverse segregation phenotype, i.e. equational division of both homologs and since this involves one completed C-CO event, the tetrad can be categorized both as C-CO or RS event. Another point to note is that MI NDJ events nearly always lead to all aneuploid spores upon one round of meiosis; however, MI PSSC, in most cases, would atleast have 50% euploid spores. Hence, crossovers at the centromeres if timely resolved or upon removal of cohesin from their vicinity (similar to that in the chromosomal arms during anaphase I), may be relatively less deleterious, and can improve the chances of obtaining euploid gametes for preventing developmental disorders arising due to trisomy in humans.

**Figure 5.**
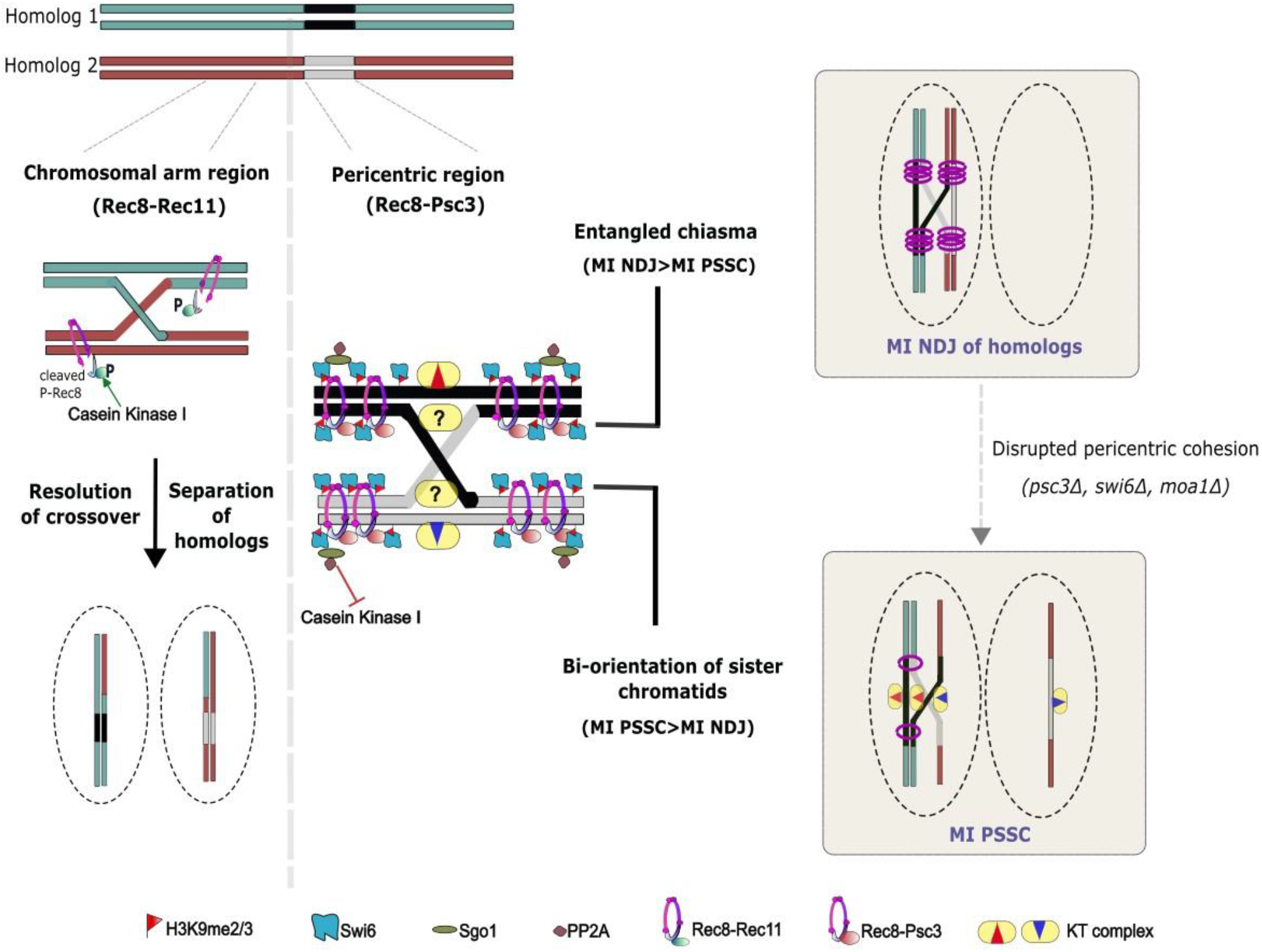
Model to explain the elevated levels of MI errors upon elevated centromeric crossovers. Occurrence of centromere-proximal crossovers (C-COs) can promote chromosomal mis-segregation via two pathways in *S. pombe*. Proteolytic cleavage of cohesins in the chromosomal arms results in resolution of chiasma during anaphase I. In contrast, the population of “protected” centromeric cohesins may hinder the proper and timely resolution of centromeric crossovers resulting in homologs being stuck or entangled with each other, unable to separate, thereby resulting in increased nondisjunction events at meiosis I. The C-COs may also disrupt monopolar attachment of the kinetochores at MI, thereby resulting in elevated propensity of biorientation of atleast one pair of sister chromatids and causing MI PSSC events. P-Rec8 stands for phosphorylated Rec8 cohesin.

## Methods

### Strains and genetic methods

The genotypes of the *S. pombe* strains generated and used in this study are described in Supplementary Table 1. Strains were generated by meiotic crosses followed by random spore analysis, as previously described^43^. Strains with temperature-sensitive alleles were grown at 25°C, while the rest were cultured at 30°C. Oligonucleotides and plasmids are described in Supplementary Tables 2 and 3, respectively. Colony PCRs were performed for genotyping.

### *S. pombe* transformation

Transformations were done using the lithium acetate method, as described previously^44^. Briefly, log phase cells at OD_600_ of 0.3-0.4 (∼1X10^7 cells) were cultured in appropriate media and after washing, resuspended in TE/LiOAc buffer (0.1 M of LiOAc in TE buffer) and incubated at 30°C for 30 min. The cells were then pelleted and resuspended in TE/LiOAc buffer containing 40% PEG4000 and 50μg boiled salmon sperm DNA (Invitrogen, 15632011), with or without ∼1 μg plasmid DNA and incubated at 30°C for 30 min. Post incubation, the cells were given a heat shock at 42°C for 20 min after adding DMSO. The cells were collected and allowed to recover in the appropriate media at 30°C for 20-22 hours, followed by selection on appropriate media. For expression of Rec10, plasmid ME46 was transformed in MP1063, MP1064, MP208, MP238, MP109, MP244, MP183, MP1011, MP1269, MP157, MP10, MP1270, MP58, MP1271, MP1383, MP1384, MP1387 and MP1388 thereby generating MP1065, MP1066, MP1012, MP1013, MP1014, MP1015, MP1016 and MP1017, MP1273, MP1274, MP1272, MP1275, MP1018, MP1276, MP1385, MP1386, MP1389 and MP1390 respectively (Suppl. Table 1). The transformants were confirmed by colony PCRs to either detect the plasmid or the correct genomic integrations.

### Cloning of *rec10* for ectopic overexpression

Using *S. pombe* genomic DNA as template, a DNA construct containing *rec10* gene flanked by BamHI (forward primer) and SmaI (reverse primer) sites was amplified using primers MO323 and MO324 (Suppl. Table 2). Plasmid ME40 and the DNA construct were digested with BamHI and SmaI and further treated with alkaline phosphatase (Takara, 2250A) to prevent self-ligation of the vector. The digested products were ligated to generate plasmid ME46 containing N-terminal *3X-myc* tagged *rec10* gene under the *Pnmt1* promoter and *Tnmt1* terminator. The recombinant plasmid ME46 was transformed into appropriate strains, confirmed by diagnostic PCRs using primers MO198, MO199, MO331 that gives a 730 bp plasmid-specific amplicon only in the transformed strains. Further, the overexpression of Rec10 was confirmed by western blotting.

### Protein expression check by immunoblotting

The expression of Rec10 from the plasmid *Pnmt1-myc-rec10* was checked by immunoblotting and cell lysates were prepared by alkaline lysis, as described previously^45^. Briefly, the strains were cultured in 10 ml NBL (-uracil) for plasmid selection and grown at 30°C up to OD_600_ of 0.6-0.8. The washed pellets were resuspended in 150μl 1.85 M NaOH and 50μl β-mercaptoethanol (7.5%) to lyse the cells followed by 15 min incubation on ice. 165μl of 50% Trichloroacetic acid was added and samples were incubated for 10 minutes on ice (neutralisation step). Samples were centrifuged at 16,800xg for 10 minutes and the pellets were suspended in a 1X Laemmli buffer (6X Laemmli buffer: 375 mM Tris-HCl (pH 6.8), 35-40% glycerol, 9% SDS, 10% v/v BME, 0.015% w/v bromophenol blue) and neutralized by Tris-HCl (pH-8.0) to make the solution alkaline. Samples were boiled to elute the protein and after centrifugation at 12,000 rpm for 1 minute, the supernatant was collected and loaded on SDS-PAGE, transferred to a PVDF membrane using transfer buffer (1X Tris Glycine, 20% Methanol) for 3 hr 15 min at a constant current of 350 mA at 4°C. The membrane was probed using 1:1000 dilutions of anti-Myc antibody (Invitrogen, MA121316) to probe for Rec10 and anti-tubulin antibody (Invitrogen, MA1-80017) to probe for tubulin as a loading control. Peroxidase AffiniPure Goat Anti-Mouse IgG (Jackson Immunoresearch, 115-035-003) was used as the secondary antibody. The blots were developed using SuperSignal West Pico PLUS Chemiluminescent Substrate (Thermo Fisher Scientific, 34580) and imaging was done using iBright CL1500 Imaging System.

### Fluorescence-based tetrad assay for chromosomal segregation

Strain generation for fluorescence-based assay was either done by genetically crossing MP83 and MP84 (and their derivatives) with the other strains of required genotypes. For *moa1*Δ strains, MP1381 and MP1382, *moa1*Δ *rec10-CD* strain MP1382 fluorophores were integrated into the *per1* locus by transformation following the protocol described above. ME29 and ME30 plasmids (Suppl. Table 3) were digested and linearised using Apa1 restriction enzyme such that the linear fragment had flanking sequences homologous to the *per1* locus. The linearised plasmid was transformed into *moa1*Δ strains with *his3-D1* mutation and His+ transformants were identified. Site specific integration was detected by PCRs using MO150, MO153 for left side integration and MO151, MO152 for right side integration, for each case giving a 1.5 kb amplicon. (Suppl. Table 2).

The parental strains were spotted on SPA media for 2-3 days at 25°C and meiosis was confirmed via asci formation. For imaging the tetrads, a small amount of the mating mixture was scraped off the SPA and mixed with a few drops of 50% glycerol on a glass slide and sealed off, to immobilize the cells. The cells were then visualized on a Leica DM6 upright epifluorescence microscope using Texas Red (TxR) filter to detect red (tdTomato) fluorescence and SBL filter to detect blue (mCerulean) fluorescence. A 63X objective was used for taking images in the brightfield (BF) and fluorescence channels. The whole slide was scanned to capture atleast 50-200 ascii per experimental replicate. ImageJ (Fiji) software was used for processing the images and analysis was done to categorize the different classes of possible tetrad profiles (Fig. 1A). The frequencies for each class of segregation profile were calculated and plotted. A minimum of 3 biological replicates were performed for each genotype and the data for individual experiments are provided in Supplementary Tables 4, 5 and 9.

### Genetic recombination assay

We measured recombination across *cen3* using prototrophic/antibiotic markers (*ura4+/hygR/kanR* and *his3+)* introduced at *chk1* and near *mid1*, respectively^36^. For *rec10ee* strains, since the *Pnmt-rec10* plasmid (ME46) has a *ura4+* marker, the *chk1::ura4+* locus in MP6, MP819, MP820 was replaced with a *kanR* marker to generate MP1269, MP1270, MP1271 respectively. The *kanR* gene was amplified from plasmid ME2 using primers MO433, MO434 (Suppl. Table 2) with flanking regions homologous to the *chk1::ura4+* locus. The amplicon was then transformed into MP6, MP819 and MP820 following the transformation protocol described above, and the transformants were identified by G418^R^ and Uracil auxotrophy. Locus specific integration was checked by performing integration check PCRs using primers MO436 and MO62 for left side integration and MO435 and MO9 for right side integration, amplifying 611 bp and 531 bp, respectively (Suppl. Table 2).

We measured recombination across *cen3* using prototrophic/antibiotic markers (*ura4+/hygR* and *his3+)* introduced at *chk1* and near *mid1*, respectively^36^. The parental strains had two different *ade6* mutations, *ade6-52* (showing pink colonies on YEA) and *ade6-M210/ade6-M26* (showing red colonies on YEA). Arm recombination was measured between the *mid1::his3+* and *ade6* interval. This interval was referred to as the *cen3-ade6* interval, since the *his3+* and *ura4+/hygR/kanR* markers flanking the centromere 3 are tightly linked. Other arm intervals used in this study are *ade6-arg1*, again on chromosome 3. Mating was induced between the two parental strains by spotting them on SPA media and keeping it for 2-3 days at 25°C. Post-asci formation (verified by microscopy), spores were harvested after glusulase and ethanol treatment, plated on YEA plate and incubated till colonies were formed. Individual colonies were randomly picked on supplemented YEA and after sufficient growth were replica-plated on different selectable media to measure growth in the absence of required supplements based on their genotypes. Recombination frequency was calculated as the percentage of recombinant colonies over the total number of colonies screened and the mean was plotted as bar graphs. For each genotype, the data was collected for atleast 3 independent parental colonies (biological repeats) as results of independent crosses. Error bars for all N=3 data represent SEM. The data for individual experiments are provided in Supplementary Tables 6, 8 and 10.

### Propidium Iodide assay

The parental strains were spotted on SPA media for 2-3 days at 25°C and meiosis was confirmed via asci formation. A small amount of the mating mixture was scraped off the SPA and resuspended in 50 ul of autoclaved distilled water. To it, 6-7ul of Propidium Iodide (1mg/mL) was added and mixed properly and cells were left for staining for 20-30 minutes. The cells were pelleted and resuspended in 10ul of 50% glycerol and put on a clean air dried glass slide and sealed off the immobilise the cells. The cells were then visualized on a Leica DM6 upright epifluorescence microscope (63X objective) using Texas Red (TxR) filter to detect red (Propidium Iodide) fluorescence. The whole slide was scanned to capture ∼100-600 asci per experimental replicate. ImageJ (Fiji) software was used for processing the images and analysis was done to categorize the different classes of tetrad profiles (Fig. 2A). The frequencies for each class of aberrant asci profile were calculated and plotted. A minimum of 3-5 biological replicates were performed for each genotype and the data for individual experiments are provided in Supplementary Table 7.

### Statistical Analysis

Data for the genetic recombination assays was analysed by using two-tailed Fisher’s exact test. Statistical significance in different error types among the various genotypes using the fluorescence-based tetrad assay was analysed by performing the two-way ANOVA test followed by Tukey’s multiple comparisons test. A few of the comparisons for the segregation subtypes among genotypes was done using Fisher’s exact test and these have been mentioned in the figure legends. All the analysis was done using GraphPad Prism 10.6.1 software for Windows, GraphPad Software, Boston, Massachusetts USA, www.graphpad.com.

## Acknowledgements

We thank Ishan Khurma, Shruti Patil, Nandeesh Sharma, Ananya Dodamani, Purva Gupta for their technical help with yeast strain generation and genotyping. We thank MN lab members for their inputs on the manuscript and helpful discussions. We acknowledge Prof. Alexander Lorenz for sharing the spore-autonomous mCerulean and tdTomato expressing plasmids (pALo196 and pALo197) and *S. pombe* strains UoA726 and UoA727. We thank Dr. Shravan K. Mishra for the expression vector pREP4X.2 and Prof. Gerald R. Smith for sharing *S. pombe* strains. S Sen acknowledges fellowship from University Grants Commission (221610216531). This study was supported by a core grant from the Department of Biotechnology, Government of India (BT/PR44595/MED/12/953/2022) and Science and Engineering Research Board (SERB) Startup Research Grant (SRG/2020/000434) to MN and IISER Pune intramural funding. We acknowledge the IISER Pune Microscopy Facility for infrastructural support.

