## Supplementary Data for "Centromere-proximal crossovers disrupt proper homologous chromosome disjunction during meiosis"

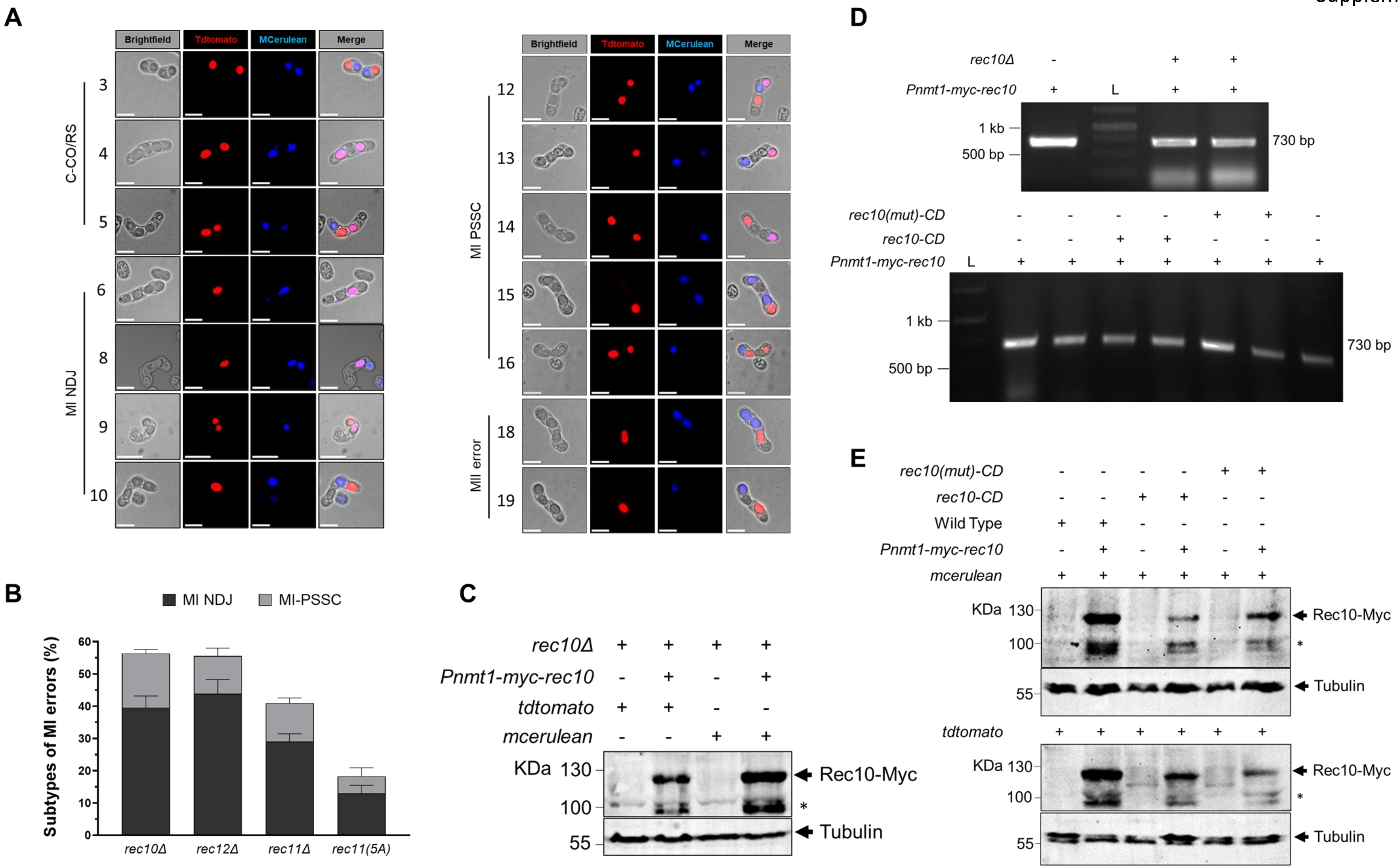
A
Brightfield
Tdtomato
MCerulean
Merge
C-CO/RS
MI NDJ
MI PSSC
MII error
3
4
5
6
8
9
10
12
13
14
15
16
18
19
B
MI NDJ
MI-PSSC
Subtypes of MI errors (%)
rec10Δ
rec12Δ
rec11Δ
rec11(5A)
C
rec10Δ
Pnmt1-myc-rec10
tdtomato
mcerulean
KDa
Rec10-Myc
Tubulin
D
rec10Δ
Pnmt1-myc-rec10
L
1 kb
500 bp
730 bp
rec10(mut)-CD
rec10-CD
E
Wild Type
mcerulean
tdtomato
KDa
Rec10-Myc
Tubulin

Supplementary Figure 2

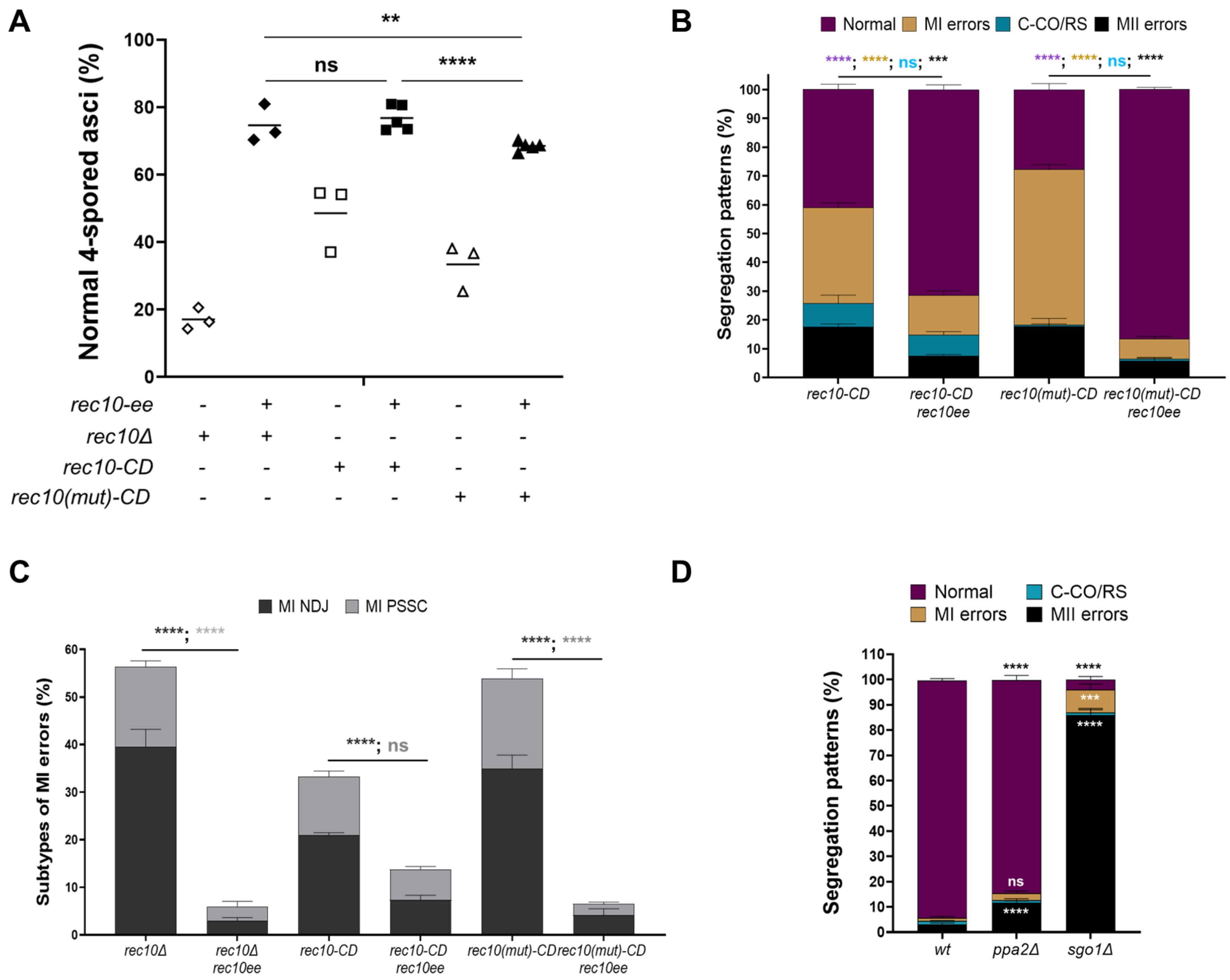

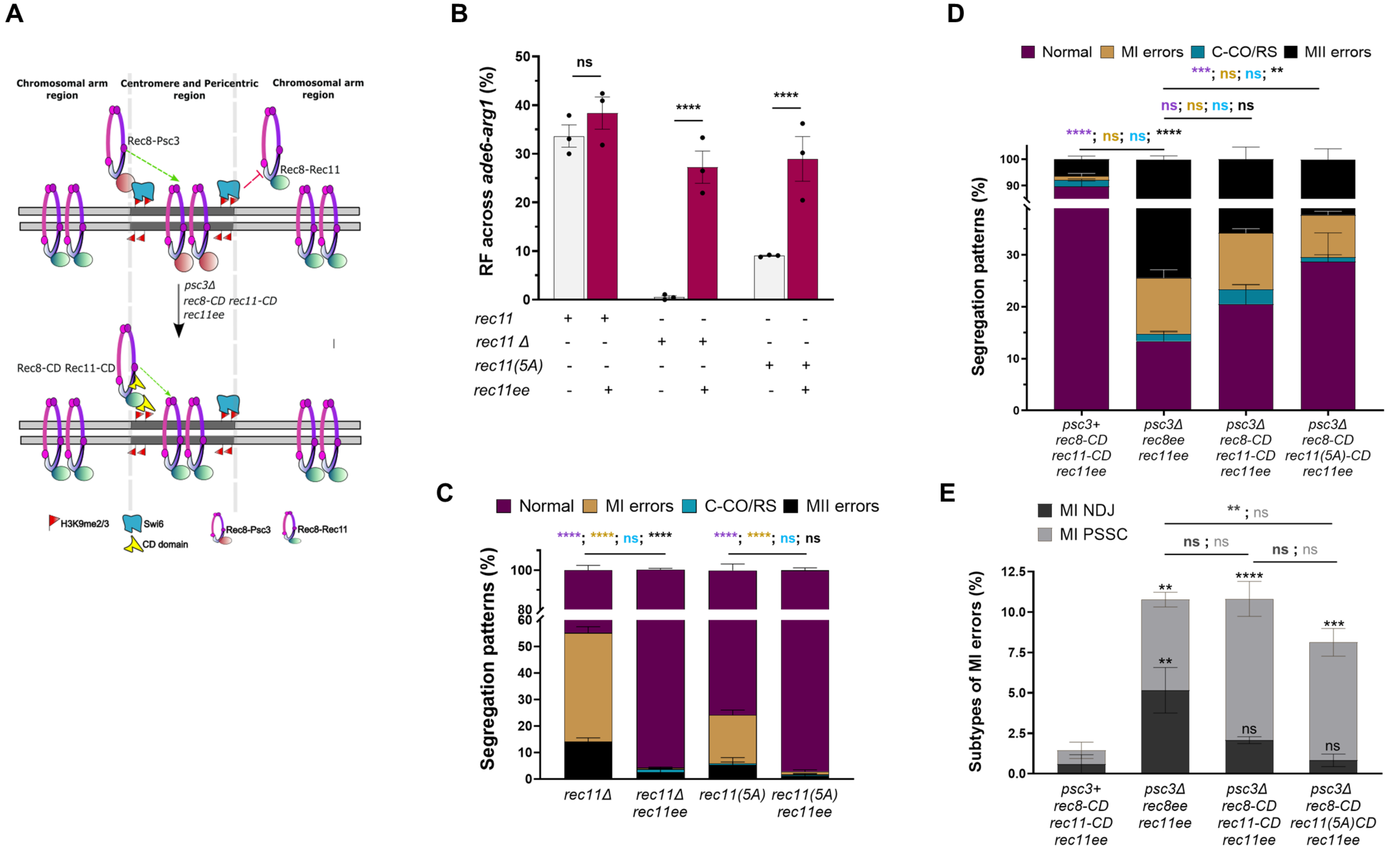
A
Chromosomal arm region
Centromere and Pericentric region
Chromosomal arm region
Rec8-Psc3
Rec8-Rec11
psc3Δ
rec8-CD rec11-CD
rec11ee
Rec8-CD Rec11-CD
H3K9me2/3
Swi6
CD domain
Rec8-Psc3
Rec8-Rec11
B
RF across ade6-arg1 (%)
ns
****
****
rec11
rec11 Δ
rec11(5A)
rec11ee
C
Normal
MI errors
C-CO/RS
MII errors
****; ****; ns; ****
****; ****; ns; ns
Segregation patterns (%)
rec11Δ
rec11Δ rec11ee
rec11(5A)
rec11(5A) rec11ee
D
Normal
MI errors
C-CO/RS
MII errors
***; ns; ns; **
ns; ns; ns; ns
****; ns; ns; ****
Segregation patterns (%)
psc3+ rec8-CD rec11-CD rec11ee
psc3Δ rec8ee rec11ee
psc3Δ rec8-CD rec11-CD rec11ee
psc3Δ rec8-CD rec11(5A)-CD rec11ee
E
MI NDJ
MI PSSC
** ; ns
ns ; ns
ns ; ns
Subtypes of MI errors (%)
psc3+ rec8-CD rec11-CD rec11ee
psc3Δ rec8ee rec11ee
psc3Δ rec8-CD rec11-CD rec11ee
psc3Δ rec8-CD rec11(5A)CD rec11ee

Supplementary Figure 4

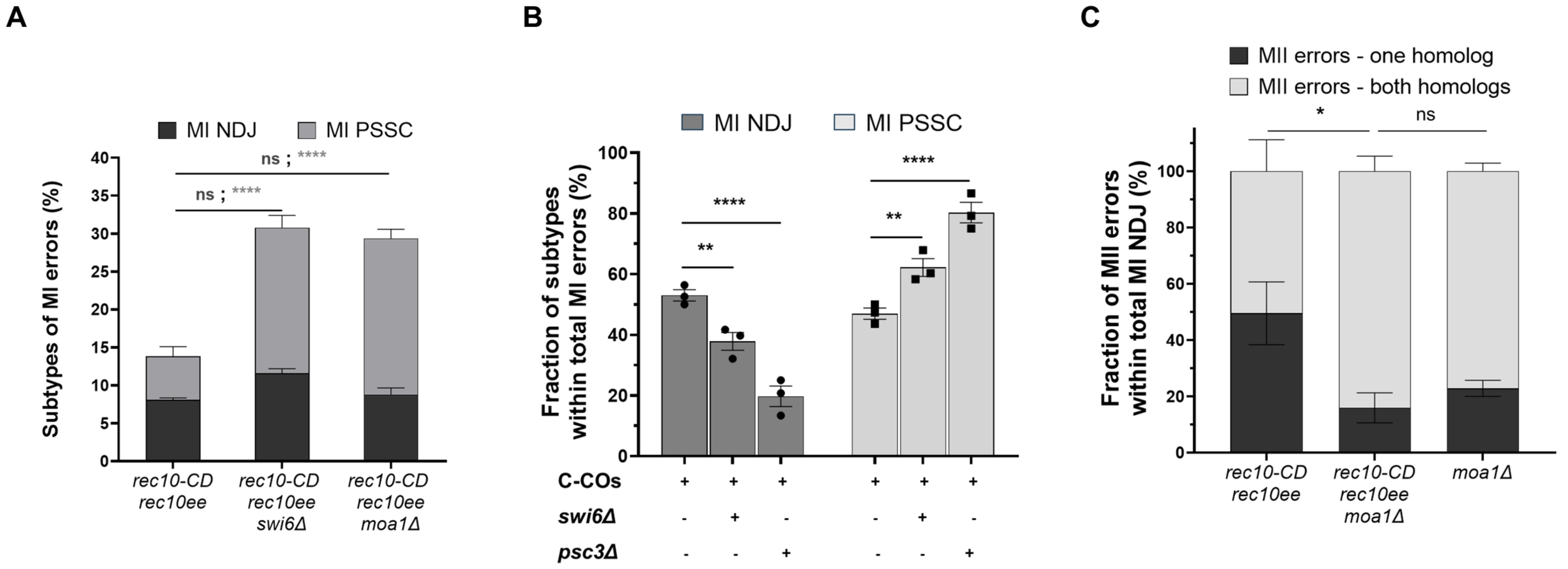

### Supplementary Figure Legends

**Supplementary Figure 1. Different segregation error types assayed during meiosis and confirmation of ectopic expression of Rec10 in the generated genotypes. A.** Representative images for the different types of normal, centromeric crossover (C-CO) or reverse segregation (RS), meiosis I non-disjunction (MI NDJ), meiosis I premature separation of sister chromatids (MI PSSC) and meiosis II (MII) error types observed. Multiple varying examples from each of these error sub-types are shown here and the numbers represent the corresponding error type as depicted in the schematic cartoon in Fig. 1A. Scale bars represent 10 µm. **B.** Bar graph showing absolute MI errors and the distribution of MI non-disjunction (NDJ) and MI premature separation of sister chromatids (PSSC) events among the total MI errors in *rec10Δ, rec12Δ, rec11Δ* and *rec11(5A)* genotypes. All data are mean ± SEM (n = 3 experiments), assaying 509, 333, 489 and 364 tetrads for *rec10Δ, rec12Δ, rec11Δ* and *rec11(5A)*, respectively. **C.** Western blot analysis for confirming the ectopic expression of the meiosis-specific protein Rec10 in both the parental *rec10Δ* and *rec10Δ rec10ee* genotypes. **D.** PCR confirmation for the presence of the plasmid overexpressing Rec10-Myc in *rec10Δ rec10ee*, *rec10ee, rec10-CD rec10ee* and *rec10(mut)-CD rec10ee* genotypes used for genetic recombination assays. **E.** Western blot analysis for confirming the ectopic expression of the meiosis-specific protein Rec10-Myc in all the parental strains of wild type, *rec10ee, rec10-CD rec10ee* and *rec10(mut)-CD rec10ee* genotypes used for segregation assays. The blots for checking Rec10 expression were probed with anti-Myc antibody. "*" denotes a non-specific band. The data for individual experiments are provided in Supplementary Table 4.

**Supplementary Figure 2. Ectopic expression of Rec10 rescues the segregation defects due to loss of arm recombination in *rec10* mutants during meiosis. A.** Rescue of spore inviability measured as the percentage of normal 4-spored asci upon Propidium Iodide staining of the meiotic tetrads in *rec10Δ, rec10Δ rec10ee, rec10-CD, rec10-CD rec10ee, rec10(mut)-CD* and *rec10(mut)-CD rec10ee* genotypes. All data are mean ± SEM (n = 3-5 experiments), assaying 430, 882, 948, 1580, 741 and 818 tetrads for *rec10Δ, rec10Δ rec10ee, rec10-CD, rec10-CD rec10ee, rec10(mut)-CD* and *rec10(mut)-CD rec10ee*, respectively. **B.** Plot showing frequency of normal, centromeric crossover (C-CO) or reverse segregation (RS), meiosis I (MI) and meiosis II (MII) segregation errors in *rec10-CD, rec10-CD rec10ee, rec10(mut)-CD* and *rec10(mut)-CD rec10ee* genotypes. **C.** Bar graph showing distribution of MI non-disjunction (NDJ) and MI premature separation of sister chromatids (PSSC) events among the total MI errors in *rec10Δ, rec10Δ rec10ee, rec10-CD, rec10-CD rec10ee, rec10(mut)-CD* and *rec10(mut)-CD rec10ee* genotypes. All data are mean ± SEM (n = 3 experiments), assaying 509, 477, 517, 479, 310 and 517 tetrads for *rec10Δ, rec10Δ rec10ee, rec10-CD, rec10-CD rec10ee, rec10(mut)-CD* and *rec10(mut)-CD rec10ee*, respectively. **D.** Plot showing frequency of normal, C-CO/RS, MI and MII segregation errors in wild type, *ppa2Δ* and *sgo1Δ* genotypes. Data are mean ± SEM (n = 3 experiments), assaying 832, 443 and 415 tetrads for wild type, *ppa2Δ* and *sgo1Δ* respectively. ****p <0.0001 (Two-way ANOVA test); ***p <0.001; **p <0.01; ns p >0.05. The data for individual experiments are provided in Supplementary Tables 5, 7 and 9.

**Supplementary Figure 3. Ectopic expression of Rec11 rescues segregation defects due to loss of arm recombination but not due to centromeric perturbations during meiosis. A.** Schematic representation of the different types of genotypes used to increase the centromeric crossover in the absence of centromere-specific cohesin Psc3 and targeting the arm cohesins Rec8 and Rec11 via chromodomain (CD) fusions. **B.** Recombinant frequency (RF) across *ade6-arg1* arm interval in *wild type, rec11ee, rec11Δ, rec11Δ rec11ee* (ectopically expressed), *rec11(5A),* and *rec11(5A) rec11*ee genotypes. Data are mean ± SEM (n = 3 experiments), assaying 390, 389, 469, 294, 672 and 411 spore colonies for *wild type, rec11ee, rec11Δ, rec11Δ rec11ee*, *rec11(5A),* and *rec11(5A) rec11ee,* respectively. ****p < 0.0001 (two-tailed Fisher's exact test); ns p >0.05. **C.** Plot showing frequency of normal, centromeric crossover (C-CO) or reverse segregation (RS), meiosis I (MI) and meiosis II (MII) segregation errors in *rec11Δ, rec11Δ rec11ee, rec11(5A)* and *rec11(5A) rec11ee* genotypes. Data are mean ± SEM (n = 3 experiments), assaying 489, 474, 364 and 417 tetrads for *rec11Δ, rec11Δ rec11ee*, *rec11(5A),* and *rec11(5A) rec11e*e, respectively. **D.** Plot showing frequency of normal, C-CO/RS, MI and MII segregation errors in *psc3+ rec8-CD rec11-CD rec11ee* (ectopically expressed), *psc3Δ rec8ee rec11ee*, *psc3Δ rec8-CD rec11-CD rec11ee* and *psc3Δ rec8-CD rec11(5A)-CD rec11ee* genotypes. **E.** Bar graph showing distribution of MI non-disjunction (NDJ) and MI premature separation of sister chromatids (PSSC) events among the total MI errors in *psc3+ rec8-CD rec11-CD rec11ee* (ectopically expressed), *psc3Δ rec8ee rec11ee*, *psc3Δ rec8-CD rec11-CD rec11ee* and *psc3Δ rec8-CD rec11(5A)-CD rec11ee* genotypes. All data are mean ± SEM (n = 3 experiments), assaying 408, 364, 477 and 647 tetrads for *psc3+ rec8-CD rec11-CD rec11ee*, *psc3Δ rec8ee rec11ee*, *psc3Δ rec8-CD rec11-CD rec11ee* and *psc3Δ rec8-CD rec11(5A)-CD rec11ee*, respectively. In all these genotypes, *rec8-CD* allele is ectopically expressed. The significance values within the bars in the graph are with respect to *psc3+ rec8-CD rec11-CD rec11ee.* For panels C-E, ****p <0.0001 (two-way ANOVA test); ***p <0.001; **p <0.01; ns p >0.05. The data for individual experiments are provided in Supplementary Tables 4, 8 and 9.

**Supplementary Figure 4. Pericentric cohesion facilitates MI nondisjunction in the presence of increased centromeric crossovers. A.** Bar graph showing distribution of MI non-disjunction (NDJ) and MI premature separation of sister chromatids (PSSC) events among the total MI errors in *rec10-CD rec10ee* and *rec10-CD rec10ee swi6Δ* genotypes. **B.** Graph comparing the fraction of NDJ and PSSC events among the total MI errors for C-COs induced genotypes, as a percentage, in the absence of Psc3 and Swi6. Data for *rec10-CD rec10ee, swi6Δ rec10-CD rec10ee* and *psc3Δ rec8-CD rec11-CD rec11ee* were used. **C.** Graph showing fraction of MI NDJ events with erroneous MII segregation involving one or both homologs in *rec10-CD rec10ee,* r*ec10-CD rec10ee moa1Δ* and *moa1Δ* genotypes. All data are mean ± SEM (n = 3 experiments), assaying 479, 713, 477, 635 and 843 tetrads for *rec10-CD rec10ee, rec10-CD rec10ee swi6Δ, psc3Δ rec8-CD rec11-CD rec11ee, rec10-CD rec10ee moa1Δ* and *moa1Δ*, respectively. ****p <0.0001 (two-way ANOVA test); **p <0.01; *p <0.05; ns p >0.05. The data for individual experiments are provided in Supplementary Tables 5 and 9.

**Supplementary Table 1 - List of *Schizosaccharomyces pombe* strains used in the study.**

| Strain # | Genotype | Source | Used in |
|---|---|---|---|
| MP83 | *h+S CEN1::his3+-PSPOG_00147-tdTomato arg3-D4 his3-D1 leu1-32 ura4-D18* | Lorenz lab | Strain generation |
| MP84 | *h-smt0 CEN1::his3+-PSPOG_00147-mCerulean arg3-D4 his3-D1 leu1-32 ura4-D18* | Lorenz lab | Strain generation |
| MP20 | *h- ade6-M26 ura4-D18 rec11-271(5A)::GFP-CD-hygR his3-D1 arg1-14* | GRS lab | Strain generation |
| MP21 | *h- ade6-52 ura4-D18 his3-D1 Padh1-rec8-3HA-ura4+ C::padh15-rec11-hygR psc3::kanR* | GRS lab | Strain generation |
| MP31 | *h90 swi6::natmx6* | GRS lab | Strain generation |
| MP30 | *h90 rec10-278::gfp-cd(W104A)-hygR chIII-imrL-tetO-ura4+ mes1::hygR ura4-D18 pNatZA31-tetR-tdtomato* | GRS lab | Strain generation |
| MP58 | *h- ade6-52 rec10-277::GFP-CD-hygR ura4-D18 his3-D1* | GRS lab | Strain generation |
| MP70 | *h+ ade6-M26 ura4-D18 his3-D1 rec8-276::Padh1-rec8-GFP-CD C::Padh15-rec11-hygR (?) psc3::kanR rec11-265::GFP-CD-hygR swi6::natR* | GRS lab | Strain generation |
| MP63 | *h+ ade6-M26 ura4-D18 his3-D1 rec11-223::ura4+ arg1-14 rec10::kanR* | GRS lab | Strain generation |
| MP64 | *h+ ade6-52 rec11-5A-GFP-rec10-BsdR ura4-D18 his3-D1 rec10::kanR* | GRS lab | Strain generation |
| MP159 | *h+ ade6-52 ura4-D18 his3-D1 leu1-32* | GRS lab | Strain generation |
| MP819 | *h+ ade6-52 ura4-D18 his3-D1 rec10-264::GFP-CD-hygR chk1::ura4+ mid1::his3+* | GRS lab | Strain generation |

| | | | |
|---|---|---|---|
| MP820 | *h+ ade6-M26 rec10-277::GFP-CD-hygR ura4-D18 his3-D1 chk1::ura4+ mid1::his3+* | GRS lab | Strain generation |
| MP1089 | *h- moa1::kanR leu1* | NBRP | Strain generation |
| MP208 | *h+ CEN1::his3+-PSPOG_00147-tdTomato his3-D1 ura4-D18* | This study | Figs. 1C, 2C and 3B |
| MP238 | *h- CEN1::his3+-PSPOG_00147-mCerulean his3-D1 ura4-D18* | This study | |
| MP1063 | *h- CEN1::his3+-PSPOG_00147-mCerulean rec10::kanR his3-D1 ura4-D18* | This study | Figs. 1C, 1E, 2D and 2E; Suppl. Fig. 1B |
| MP1064 | *h+ CEN1::his3+-PSPOG_00147-tdTomato rec10::kanR his3-D1 ura4-D18 arg1-14* | This study | |
| MP751 | *h- ade6-M26 ura4-D18 rec12-169::kanR CEN1::his3+-PSPOG_00147-tdTomato* | This study | Figs. 1C, 2D and 2E; Suppl. Fig. 1B |
| MP769 | *h+ his3-D1 ura4-D18 rec12-169::kanR CEN1::his3+-PSPOG_00147-mCerulean* | This study | |
| MP330 | *h +ade6-52 ura4-D18 rec11::kanR his3-D1CEN1::his3+-PSPOG_00147-tdTomato* | This study | Fig. 1C, Suppl. Figs. 1B and 3C |
| MP331 | *h- ade6-52 ura4-D18 rec11::kanR his3-D1 CEN1::his3+-PSPOG_00147-mCerulean leu1-32* | This study | |
| MP566 | *h- ade6-52 ura4-D18 rec11-270(5A) CEN1::his3+-PSPOG_00147-mCerulean arg1-14* | This study | Fig. 1C; Suppl. Figs. 1B and 3C |
| MP568 | *h+ ade6-52 ura4-D18 rec11-270(5A) CEN1::his3+-PSPOG_00147-tdTomato arg1-14* | This study | |
| MP1065 | *h- CEN1::his3+-PSPOG_00147-mCerulean rec10::kanR his3-D1 ura4-D18 ura4-pnmt1-myc-rec10 (ME46)* | This study | Fig. 1E; |

| | | | |
|---|---|---|---|
| MP1066 | *h+ CEN1::his3+-PSPOG_00147-tdTomato rec10::kanR his3-D1 ura4-D18 arg1-14 ura4-pnmt1-myc-rec10 (ME46)* | This study | Suppl. Fig. 1C |
| MP244 | *h- rec10-264::GFP-CD-hygR CEN1::his3+PSPOG_00147-mCerulean ura4-D18 his3-D1* | This study | Suppl. Figs. 1E, 2B and 2C |
| MP109 | *h+ CEN1::his+-PSOG_00147-tdtomato leu1-32 ura4-D18 rec10-264::GFP-CD-HygR his3-D1* | This study | |
| MP183 | *h+ rec10-277::GFP-CD-hygR CEN1::his3+-PSPOG_00147-tdTomato ura4-D18 leu1-32* | This study | Suppl. Figs. 1E, 2B and 2C |
| MP1011 | *h- rec10-277::GFP-CD-hygR CEN1::his3+-PSPOG_00147- mCerulean ura4-D18* | This study | |
| MP213 | *h+ rec10-278::gfp-cd(W104A)-hygR CEN1::his3+-PSPOG_00147-tdTomato ura4-D18* | This study | Fig. 2C |
| MP217 | *h- rec10-278::GFP-CD(W104A)-hygR CEN1::his3+-PSPOG_00147-mCerulean ura4-D18 his3-D1* | This study | |
| MP1012 | *h- CEN1::his3+-PSPOG_00147-tdTomato his3-D1 ura4-D18 ura4+-Pnmt1-myc-rec10 (ME46)* | This study | Fig. 2C; Suppl. Fig. 1E |
| MP1013 | *h+ CEN1::his3+-PSPOG_00147-mCerulean ura4-D18 his3-D1 ura4+-Pnmt1-myc-rec10 (ME46)* | This study | |
| MP1014 | *h- rec10-264::GFP-CD-HygR CEN1::his3+PSPOG_00147-mCerulean ura4-D18 his3-D1 ura4+-Pnmt1-myc-rec10 (ME46)* | This study | Figs. 2C, 2D, 2E, 4B, 4C, 4D, 4E and 4F; Suppl. Figs. 1E, 2B, 2C, 4A, 4B and 4C |
| MP1015 | *h+ CEN1::his+-PSOG_00147-tdtomato leu1-32 ura4-D18 rec10-264::GFP-CD-hygR his3-D1 ura4+-Pnmt1-myc-rec10 (ME46)* | This study | |
| MP1016 | *h+ rec10-277::GFP-CD-HgR CEN1::his3+-PSPOG_00147-tdTomato ura4-D18 leu1-32 ura4+-Pnmt1-myc-rec10 (ME46)* | This study | Fig. 2C; |

| | | | |
|---|---|---|---|
| MP1017 | *h- rec10-277::GFP-CD-hygR CEN1::his3+-PSPOG_00147- mCerulean ura4-D18 ura4+-Pnmt1-myc-rec10 (ME46)* | This study | Suppl. Figs. 2B and 2C |
| MP578 | *h- ade6-52 ura4-D18 CEN1::his3+-PSPOG_00147-mCerulean leu1-32 rec11::kanR C::Padh15-rec11-hygR* | This study | Suppl. Fig. 3C |
| MP607 | *h- ura4-D18 CEN1::his3+-PSPOG_00147-Tdtomato arg3-D4 rec11::kanR C::Padh15-rec11-hygR* | This study | |
| MP777 | *h- ade6-52 ura4-D18 rec11-270(5A) leu1-32 CEN1::his3+-PSPOG_00147-mCerulean C::Padh15-rec11-hygR* | This study | Suppl. Fig. 3C |
| MP779 | *h+ ade6-52 ura4-D18 CEN1::His3+_PSPOG_OO147_tdtomato rec11-270(5A) C::Padh15-rec11-hygR arg3Δ* | This study | |
| MP368 | *h- CEN1::his3+-PSPOG_00147-mCerulean his3-D1 ura4-D18 Padh1-rec8-3HA-ura4+ C::Padh15-rec11-hygR psc3::kanR* | This study | Figs. 3B and 4A; Suppl. Figs. 3D and 3E |
| MP387 | *h+ CEN1::his3+-PSPOG_00147-tdTomato his3-D1 ura4-D18 Padh1-rec8-3HA-ura4+ C::Padh15-rec11-hygR psc3::kanR* | This study | |
| MP288 | *h+ CEN1::his3+-PSPOG_00147-mCerulean his3-D1 ura4-D18 swi6::natR* | This study | Figs. 3B and 4B |
| MP366 | *h- CEN1::his3+-PSPOG_00147-tdTomato his3-D1 ura4-D18 swi6::natR* | This study | |
| MP745 | *h- ura4-D18 his3-D1 sgo1::kanR CEN1::his3+-PSPOG_00147-tdTomato* | This study | Fig. 3B; Suppl. Fig. 2D |
| MP746 | *h+ ade6-52 ura4-D18 his3-D1 sgo1::kanR CEN1::his3+-PSPOG_00147-mCerulean* | This study | |
| MP754 | *h- his3-D1 ppa2::ura4+ ura4-D18 CEN1::his3+-PSPOG_00147-tdTomato* | This study | Suppl. Fig. 2D |

| | | | |
|---|---|---|---|
| MP770 | *h+ his3-D1 ppa2::ura4+ ura4-D18 CEN1::his3+-PSPOG_00147-mCerulean leu1-32* | This study | |
| MP772 | *h+ ade6-M26 ura4-D18 his3-D1 lys4-95 clr4::kanR CEN1::his3+-PSPOG_00147-tdTomato* | This study | Fig. 3B |
| MP749 | *h- ade6-52 ura4-D18 his3-D1 clr4::kanR CEN1::his3+-PSPOG_00147-mCerulean* | This study | |
| MP713 | *h+ ade6-52 ura4 -D18 chp2::natR CEN1:his3+-PSPOG_00147Mcerulean* | This study | Fig. 3B |
| MP756 | *h- ade6-52 ura4-D18 his3-D1 chp2::natR CEN1::his3+-PSPOG_00147-tdTomato* | This study | |
| MP718 | *h+ ade6-M26 CEN1::his3+-PSPOG_00147-tdTomato his3-D1 ura4-D18 rec8-276::Padh1-rec8-GFP-CD C::Padh15-rec11-hygR rec11-265::GFP-CD-hygR* | This study | Suppl. Figs. 3D and 3E |
| MP719 | *h- ade6-M26 CEN1::his3+-PSPOG_00147-mCerulean his3-D1 ura4-D18 rec8-276::Padh1-rec8-GFP-CD C::Padh15-rec11-hygR (?) rec11-265::GFP-CD-hygR* | This study | |
| MP720 | *h+ ade6-M26 CEN1::his3+-PSPOG_00147-mCerulean his3-D1 ura4-D18 rec8-276::Padh1-rec8-GFP-CD C::Padh15-rec11-hygR rec11-265::GFP-CD-hygR psc3::kanR* | This study | Fig. 4A; Suppl. Figs. 3D, 3E and 4B |
| MP721 | *h- ade6-M26 CEN1::his3+-PSPOG_00147-tdTomato his3-D1 ura4-D18 rec8-276::Padh1-rec8-GFP-CD C::Padh15-rec11-hygR rec11-265::GFP-CD-hygR psc3::kanR* | This study | |
| MP722 | *h+ his3-D1 ura4-D18 CEN1::his3+-PSPOG_00147-mCerulean rec11-271(5A)::GFP-CD-hygR rec8-276::Padh1-rec8-GFP-CD c::Padh15-rec11-hygR psc3::kanR* | This study | Fig. 4A; Suppl. Figs. 3D and 3E |
| MP723 | *h- his3-D1 ura4-D18 CEN1::his3+-PSPOG_00147-tdTomato rec11-271(5A)::GFP-CD-hygR rec8-276::Padh1-rec8-GFP-CD c::Padh15-rec11-hygR psc3::kanR* | This study | |

| | | | |
|---|---|---|---|
| MP1183 | *h- rec10-264::GFP-CD-hygR CEN1::his+-PSOG_00147-tdtomato ura4-D18 his3-D1 ura4+-Pnmt1-myc-rec10 (ME46) swi6::natR* | This study | Figs. 4B and 4C; Suppl. Figs. 1D, 4A and 4B |
| MP1184 | *h+ CEN1::his3+PSPOG_00147-mCerulean leu1-32 ura4-D18 rec10-264::GFP-CD-hygR his3-D1 ura4+-Pnmt1-myc-rec10 (ME46) swi6::natR* | This study | |
| MP1382 | *h- moa1::kanR his3-D1 CEN1::his3+-PSPOG_00147-tdTomato leu1-32* | This study | Figs. 4B, 4D, 4E and 4F; Suppl. Fig. 4C |
| MP1381 | *h+ moa1::kanR his3-D1 CEN1::his3+-PSPOG_00147-mCerulean ura4-D18 leu1-32* | This study | |
| MP1385 | *h+ rec10-264::GFP-CD-hygR moa1::kanR ura4-D18 his3-D1 CEN1::his3+-PSPOG_00147-tdTomato ura4+-Pnmt1-myc-rec10 (ME46) leu1-32* | This study | Figs. 4B, 4D, 4E and 4F; Suppl. Figs. 1D, 4A and 4C |
| MP1386 | *h- rec10-264::GFP-CD-hygR moa1::kanR ura4-D18 his3-D1 CEN1::his3+-PSPOG_00147-mCerulean ura4+-Pnmt1-myc-rec10 (ME46) leu1-32* | This study | |
| MP6 | *h- ade6-52 ura4-D18 his3-D1 mid1::his3+ chk1::ura4+* | GRS Lab | Figs. 1F, 1G and 3C |
| MP8 | *h-ade6-52 ura4-D18 his3-D1 arg1-14* | GRS Lab | |
| MP157 | *h+ ade6-M26 ura4-D18 his3-D1* | GRS Lab | |
| MP1269 | *h- ade6-52 ura4-D18 his3-D1 mid1::his3+ chk1::kanR* | This study | |
| MP1273 | *h+ ade6-M26 ura4-D18 his3-D1 ura4+-Pnmt1-myc-rec10 (ME46)* | This study | Figs. 1F, 1G and 2B; Suppl. Fig. 2A |
| MP1274 | *h- ade6-52 ura4-D18 his3-D1 mid1::his3+ chk1::kanR ura4+-Pnmt1-myc-rec10 (ME46)* | This study | |
| MP1388 | *h- ade6-52 ura4-D18 his3-D1 rec10::KanR* | This study | Fig. 1D; Suppl. Fig. 2A |
| MP1387 | *h+ ade6-M26 ura4-D18 his3-D1 chk1::hygR mid1::his3+ rec10::KanR* | This study | |

| | | | |
|---|---|---|---|
| MP1390 | *h- ade6-52 ura4-D18 his3-D1 rec10::kanR ura4+-Pnmt1-myc-rec10 (ME46)* | This study | Figs. 1D and 2B; Suppl. Fig. 2A |
| MP1389 | *h+ ade6-M26 ura4-D18 his3-D1 chk1::hygR mid1::his3+ rec10::kanR ura4+-Pnmt1-myc-rec10 (ME46)* | This study | |
| MP10 | *h- ade6-M26 ura4-D18 his3-D1 rec10-264::GFP-CD-hygR* | GRS Lab | Fig. 1F and 1G; Suppl. Fig. 2A |
| MP1270 | *h+ ade6-52 ura4-D18 his3-D1 rec10-264::GFP-CD-hygR chk1::kanR mid1::his3+* | This study | |
| MP1272 | *h- ade6-M26 ura4-D18 his3-D1 rec10-264::GFP-CD-hygR ura4+-Pnmt1-myc-rec10 (ME46)* | This study | Figs. 1F, 1G and 2B; Suppl. Fig. 2A |
| MP1275 | *h+ ade6-52 ura4-D18 his3-D1 rec10-264::GFP-CD-hygR chk1::kanR mid1::his3+ ura4+-Pnmt1-myc-rec10 (ME46)* | This study | |
| MP58 | *h- ade6-52 rec10-277::GFP-CD-hygR ura4-D18 his3-D1* | GRS Lab | Fig. 1F and 1G; Suppl. Fig. 2A |
| MP1271 | *h+ ade6-M26 rec10-277::GFP-CD-hygR ura4-D18 his3-D1 chk1::kanR mid1::his3+* | This study | |
| MP1018 | *h- ade6-52 rec10-277::GFP-CD-hygR ura4-D18 his3-D1 ura4+-Pnmt1-myc-rec10 (ME46)* | This study | Figs. 1F, 1G and 2B; Suppl. Fig. 2A |
| MP1276 | *ade6-M26 rec10-277::GFP-CD-hygR ura4-D18 his3-D1 chk1::kanR mid1::his3+ ura4+-Pnmt1-myc-rec10 (ME46)* | This study | |
| MP498 | *h- ade6-52 arg1-1 ura4-D18 c::Padh15-rec11-hygR* | This study | Suppl. Fig. 3B |
| MP499 | *h+ ade6-M210 ura4-D18 c::Padh15-rec11-hygR* | This study | |
| MP450 | *h- ade6-52 ura4-D18 arg1-14 rec11::kanR* | This study | Suppl. Fig. 3B |
| MP453 | *h+ ade6-M210 his3-D1 ura4-D18 rec11::kanR* | This study | |
| MP782 | *h+ ade6-M210 c::padh15-rec11-hygR rec11::kanR* | This study | |

| | | | |
|---|---|---|---|
| MP797 | *h- ade6-52 ura4-D18 arg1-14 c::padh15-rec11-hygR rec11::kanR* | This study | Suppl. Fig. 3B |
| MP520 | *h+ ade6-52 rec11-270(5A) arg1-14* | GRS lab | Suppl. Fig. 3B |
| MP521 | *h- ade6-M26 rec11-270(5A)* | GRS lab | |
| MP796 | *h+ ade6-M26 Ura4-D18 his3-D1 rec11-270(5A) c::Padh15-rec11-hygR* | This study | Suppl. Fig. 3B |
| MP883 | *h- ade6-52 CEN1::His3+_PSPOG_OO147_mcerulean rec11-270(5A) c::Padh15-rec11-hygR arg1-14* | This study | |
| MP641 | *h- ade6-M26 ura4-D18 chk::ura4+ mid1::his3+ sgo1::kanR his3-D1* | GRS lab | Fig. 3C |
| MP642 | *h+ ade6-52 ura4-D18 his3-D1 lys4-95 sgo1::kanR* | GRS lab | |
| MP78 | *h+ ade6-52 ura4-D18 his3-D1 clr4::kanR* | GRS lab | Fig. 3C |
| MP639 | *h- ade6-M26 ura4-D18 his3-D1 lys+ chk1::ura4+ mid1::his3+ clr4::kanR* | GRS lab | |

**Supplementary Table 2** - List of oligonucleotides used in the study.

| Oligomer # | Sequence | Purpose |
|---|---|---|
| MO1 | AGAAGAGAGAGTAGTTGAAG | Mating type |
| MO2 | ACGGTAGTCATCGGTCTTCC | Mating type |
| MO3 | TACGTTCAGTAGACGTAGTG | Mating type |
| MO25 | GCGAACTGTAAACAATGGTGC | *his3-D1* |
| MO26 | GCGTAAAACTCGGATTTATGCTAGG | *his3-D1* |
| MO22 | GGTGTTGGTGGATCCAACTGGC | *ura4-D18* |
| MO41 | CATGAAGAATTGGTTATCCGTGTGC | *ura4-D18* |
| MO42 | GTTTCAGAAGAACATTACTCGTCGTG | *ura4-D18* |
| MO40 | AATCGTGCTAGAGATCCTGG | *hygR* Fwd |
| MO45 | CCCCAATGTCAAGCACTTCC | *rec11* 3'-UTR Rev |
| MO198 | AGAATTATTGTTGTCTCTTTATGG | *Pnmt1* Fwd |
| MO199 | AATATTCAAAACGATTTAATGCC | *Tnmt1* Rev |
| MO331 | GTCAAGCTCGAGGAAATACC | *rec10* ORF Fwd |
| MO323 | CGTAGTGGATCCGCTGCTGCTATGGACATATTTTCTGAGTTTCTC | *rec10* cloning in ME40; with BamHI |
| MO324 | ACTACGCCCGGGCTACTTAAACATCAAACTGTCAATTTG | *rec10* cloning in ME40; with SmaI |
| MO433 | TTTCCTTACCATATTGGTAGAGAAATAGGTACTGGGGCTTTTGCTTCGTATGAATTTTGATTTGAAAAATACTATATACTAGATCTGTTTAGCTTGCCTC | *chk1_kanR* Fwd |
| MO434 | ACTCCTTTTGGTCATTTTTTTTAAAAGCTGGCTGTGTCATTGAAAAACCGCTATCGCTAATTGAAAGAGTTAGAAATATGGGATGGCGGCGTTAGTATCG | *chk1_kanR*_Rev |
| MO62 | GATTGCCCGACATTATCGCG | *kanR* ORF Rev |
| MO436 | CTTTCCCTTTACACCAAAGAATGG | *chk1* 5'UTR Fwd |
| MO9 | CCTCGGTGAGTTTTCTCC | *kanR* ORF Fwd |
| MO435 | ACAAAACTGACTCAATGAGGG | *chk1* ORF Rev |
| MO150 | ATCGTCTTCCGTCTTGCTATG | *per1 cen1L* Fwd |
| MO153 | CTCCCGATATGTCTGCCTTAAC | *his3* Rev |
| MO151 | CTCCGATCGTTGTCAGAAGTAAG | *ampR* Fwd |
| MO152 | GGTCCTACAATACACGCAGATAAG | *cen1R* Rev |

**Supplementary Table 3** - List of plasmids used in the study.

| Plasmid | Genotype | Source |
|---|---|---|
| ME29 (pALo196) | PSPOG_00147- tdTomato-TPGK1(Skud) (targeting to CEN1) | Prof. A. Lorenz (UoA) |
| ME30 (pALo197) | PSPOG_00147- mCerulean-TPGK1(Sbay) (targeting to CEN1) | Prof. A. Lorenz (UoA) |
| ME40 (pREP4X.2) | 3XMYC-MCS-EGFP (pnmt1 promoter, ura4+ selection) | Dr. S. K. Mishra (IISERM) |
| ME46 | Rec10 ORF cloned into ME40 (EGFP removed, 3XMYC used as N-terminal tag for Rec10) | This study |
| ME2 | pFA6a-GFP-KanMX6 | Prof. G. R. Smith (Fred Hutch) |

**Supplementary Table 4.** Analysis of segregation defects at c*en1* during meiosis, related to Figure 1, and Supplementary Figure 1.

| | **Genotype** | **Strains crossed** | **Trials** | **Total tested** | **Segregation pattern** | | | | |
|---|---|---|---|---|---|---|---|---|---|
| | | | | | Normal | C-CO/RS | Meiosis I | | Meiosis II |
| | | | | | | | NDJ | PSSC | |
| 1 | Wild Type | MP208 x MP238 | 1 | 144 | 136 | 2 | 0 | 1 | 5 |
| | | | 2 | 177 | 168 | 1 | 0 | 0 | 8 |
| | | | 3 | 57 | 55 | 0 | 0 | 2 | 0 |
| | | | 4 | 136 | 134 | 0 | 0 | 0 | 2 |
| | | | 5 | 258 | 244 | 1 | 1 | 1 | 11 |
| | | | 6 | 62 | 62 | 0 | 0 | 0 | 0 |
| 2 | *rec12Δ* | MP751 x MP769 | 1 | 59 | 14 | 3 | 23 | 4 | 15 |
| | | | 2 | 160 | 44 | 0 | 64 | 23 | 29 |
| | | | 3 | 114 | 19 | 1 | 60 | 16 | 18 |
| 3 | *rec11Δ* | MP330 x MP331 | 1 | 148 | 63 | 0 | 38 | 22 | 25 |
| | | | 2 | 237 | 118 | 0 | 66 | 22 | 31 |
| | | | 3 | 104 | 44 | 0 | 35 | 12 | 13 |
| 4 | *rec11(5A)* | MP566 x MP568 | 1 | 179 | 124 | 3 | 17 | 19 | 16 |
| | | | 2 | 96 | 75 | 0 | 11 | 3 | 7 |
| | | | 3 | 89 | 71 | 0 | 16 | 2 | 0 |
| 5 | *rec11Δ rec11ee* | MP578 x MP607 | 1 | 109 | 103 | 1 | 0 | 0 | 5 |
| | | | 2 | 215 | 205 | 3 | 1 | 1 | 5 |
| | | | 3 | 150 | 146 | 2 | 0 | 1 | 1 |
| 6 | *rec11(5A) rec11ee* | MP777 x MP779 | 1 | 150 | 145 | 2 | 0 | 1 | 2 |
| | | | 2 | 82 | 78 | 0 | 1 | 1 | 2 |
| | | | 3 | 185 | 184 | 0 | 0 | 1 | 0 |

**Supplementary Table 5.** Analysis of segregation defects at c*en1* during meiosis, related to Figures 1, 2, 4 and Supplementary Figures 2 and 4.

| | Genotype | Strains crossed | Trials | Total tested | Segregation pattern (# of tetrads) | | | | |
|---|---|---|---|---|---|---|---|---|---|
| | | | | | Normal | C-CO/RS | Meiosis I | | Meiosis II |
| | | | | | | | NDJ | PSSC | |
| 1 | Wild type | MP208 x MP238 | 1 | 143 | 139 | 3 | 0 | 0 | 1 |
| | | | 2 | 163 | 155 | 0 | 1 | 1 | 6 |
| | | | 3 | 82 | 80 | 0 | 0 | 1 | 1 |
| | | | 4 | 133 | 129 | 1 | 0 | 0 | 3 |
| | | | 5 | 184 | 171 | 3 | 1 | 1 | 8 |
| | | | 6 | 185 | 175 | 0 | 1 | 1 | 8 |
| 2 | *rec10*ee | MP1012 x MP1013 | 1 | 94 | 88 | 1 | 0 | 1 | 4 |
| | | | 2 | 111 | 106 | 0 | 0 | 0 | 5 |
| | | | 3 | 162 | 153 | 0 | 3 | 2 | 4 |
| 3 | *rec10-CD* | MP244 x MP109 | 1 | 195 | 76 | 17 | 41 | 23 | 38 |
| | | | 2 | 75 | 30 | 9 | 15 | 8 | 13 |
| | | | 3 | 247 | 110 | 7 | 54 | 36 | 40 |
| 4 | *rec10(mut)-CD* | MP183 x MP1011 | 1 | 91 | 22 | 0 | 27 | 21 | 21 |
| | | | 2 | 111 | 35 | 1 | 39 | 18 | 18 |
| | | | 3 | 108 | 30 | 1 | 43 | 19 | 15 |
| 5 | *rec10-CD(mut)* | MP213 x MP217 | 1 | 236 | 216 | 3 | 3 | 3 | 11 |
| | | | 2 | 128 | 115 | 0 | 6 | 4 | 3 |
| | | | 3 | 196 | 186 | 5 | 1 | 1 | 3 |
| 6 | *rec10Δ* | MP1063 x MP1064 | 1 | 136 | 44 | 1 | 44 | 26 | 21 |
| | | | 2 | 212 | 51 | 0 | 91 | 32 | 38 |
| | | | 3 | 161 | 37 | 1 | 70 | 26 | 27 |
| 7 | *rec10Δ rec10ee* | MP1065 x MP1066 | 1 | 196 | 171 | 1 | 6 | 6 | 12 |
| | | | 2 | 99 | 91 | 2 | 4 | 1 | 1 |

| | | | | | | | | | |
|---|---|---|---|---|---|---|---|---|---|
| | | | 3 | 182 | 153 | 3 | 3 | 9 | 14 |
| 8 | *rec10-CD rec10ee* | MP1014 x MP1015 | 1 | 119 | 81 | 10 | 10 | 9 | 9 |
| | | | 2 | 95 | 69 | 8 | 5 | 5 | 8 |
| | | | 3 | 265 | 195 | 13 | 22 | 17 | 18 |
| 9 | *rec10(mut)-CD rec10ee* | MP1016 x MP1017 | 1 | 116 | 99 | 2 | 8 | 2 | 5 |
| | | | 2 | 125 | 109 | 0 | 3 | 3 | 10 |
| | | | 3 | 276 | 242 | 2 | 8 | 10 | 14 |
| 10 | *rec10-CD rec10ee swi6Δ* | MP1183 x MP1184 | 1 | 201 | 50 | 5 | 23 | 35 | 88 |
| | | | 2 | 276 | 68 | 7 | 35 | 49 | 117 |
| | | | 3 | 236 | 54 | 6 | 25 | 53 | 98 |
| 11 | *rec10-CD rec10ee moa1Δ* | MP1385 x MP1386 | 1 | 149 | 18 | 4 | 13 | 28 | 86 |
| | | | 2 | 290 | 41 | 12 | 27 | 66 | 144 |
| | | | 3 | 196 | 23 | 3 | 14 | 40 | 116 |

**Supplementary Table 6.** Analysis of recombination between markers flanking *cen3* and at arm interval *cen3-ade6* during meiosis, related to Figure 1.

| | Genotype | Strains crossed | No. of crosses | Total tested | *cen3* recombination (RF) | | | Arm recombination (RF) | |
|---|---|---|---|---|---|---|---|---|---|
| | | | | | $Hyg^S$ /$Kan^S$ $His^+$ | $Hyg^R$ /$Kan^R$ $His^o$ | *cen3* RF (%) | No. of recombinants | *cen3-ade6* RF (%) |
| 1 | Wild type | MP1269 x MP157 | 3 | 410 | 0 | 0 | - | 58 | 14.1 |
| 2 | *rec10Δ* | MP1388 x MP1387 | 3 | 408 | 0 | 0 | - | 5 | 1.2 |
| 3 | *rec10Δ rec10ee* | MP1389 x MP1390 | 3 | 408 | 0 | 1 | 0.25 | 50 | 12.3 |
| 4 | *rec10ee* | MP1272 x MP1274 | 3 | 412 | 0 | 0 | - | 50 | 12.1 |
| 5 | *rec10-CD* | MP10 x MP1270 | 3 | 408 | 6 | 4 | 2.5 | 7 | 1.7 |
| 6 | *rec10-CD rec10ee* | MP1272 x MP1275 | 9 | 1122 | 11 | 14 | 2.2 | 41 | 3.7 |
| 7 | *rec10(mut)-CD* | MP58 x MP1271 | 3 | 401 | 1 | 1 | 0.5 | 0 | - |
| 8 | *rec10(mut)-CD rec10ee* | MP1276 x MP1018 | 6 | 631 | 2 | 1 | 0.48 | 41 | 6.5 |

**Supplementary Table 7.** Analysis of Propidium Iodide (PI) stained upon completion of meiosis, related to Figure 2 and Supplementary Figure 2.

| | Genotype | Strains crossed | Trials | Total tested | Normal spores (N) | Abnormal spores (AS) | | | | N (%) | AS (%) |
|---|---|---|---|---|---|---|---|---|---|---|---|
| | | | | | | 1 | 2 | 3 | 4 | | |
| 1 | *rec10ee* | MP1272 x MP1274 | 1 | 137 | 122 | 12 | 2 | 1 | 0 | 89.05 | 10.95 |
| | | | 2 | 170 | 133 | 25 | 11 | 1 | 0 | 78.24 | 21.77 |
| | | | 3 | 139 | 111 | 19 | 7 | 0 | 2 | 79.85 | 20.15 |
| | | | 4 | 337 | 285 | 32 | 9 | 4 | 7 | 84.57 | 15.43 |
| | | | 5 | 276 | 246 | 18 | 8 | 2 | 2 | 89.13 | 10.87 |
| 2 | *rec10-CD* | MP10 x MP819 | 1 | 623 | 231 | 144 | 167 | 50 | 31 | 37.08 | 62.92 |
| | | | 2 | 229 | 125 | 54 | 40 | 6 | 4 | 54.59 | 45.42 |
| | | | 3 | 96 | 52 | 21 | 22 | 1 | 0 | 54.17 | 45.83 |
| 3 | *rec10-CD rec10ee* | MP1272 x MP1275 | 1 | 344 | 252 | 52 | 31 | 8 | 1 | 73.26 | 26.74 |
| | | | 2 | 294 | 238 | 24 | 23 | 6 | 3 | 80.95 | 19.05 |
| | | | 3 | 397 | 300 | 42 | 38 | 9 | 8 | 75.56 | 24.43 |
| | | | 4 | 328 | 241 | 40 | 29 | 3 | 15 | 73.48 | 26.52 |
| | | | 5 | 217 | 175 | 27 | 11 | 1 | 3 | 80.65 | 19.35 |
| 4 | *rec10(mut)-CD* | MP58 x MP1271 | 1 | 273 | 104 | 54 | 91 | 13 | 11 | 38.10 | 61.91 |
| | | | 2 | 346 | 127 | 82 | 106 | 10 | 21 | 36.71 | 63.30 |
| | | | 3 | 122 | 31 | 24 | 52 | 7 | 8 | 25.41 | 74.59 |
| 5 | *rec10(mut)-CD rec10ee* | MP1276 x MP1018 | 1 | 88 | 60 | 14 | 11 | 2 | 1 | 68.18 | 31.82 |
| | | | 2 | 173 | 119 | 26 | 19 | 4 | 5 | 68.79 | 31.21 |
| | | | 3 | 164 | 109 | 22 | 22 | 7 | 4 | 66.46 | 33.54 |
| | | | 4 | 182 | 128 | 26 | 21 | 6 | 1 | 70.33 | 29.67 |
| | | | 5 | 211 | 145 | 29 | 27 | 6 | 4 | 68.72 | 31.28 |
| 6 | *rec10 Δ* | MP1387 x MP1388 | 1 | 104 | 17 | 12 | 43 | 7 | 25 | 16.35 | 83.65 |
| | | | 2 | 165 | 34 | 13 | 72 | 12 | 34 | 20.61 | 79.39 |

| | | | | | | | | | | | |
|---|---|---|---|---|---|---|---|---|---|---|---|
| | | | 3 | 161 | 23 | 18 | 57 | 21 | 42 | 14.29 | 85.71 |
| 7 | *rec10* Δ *rec10ee* | MP1389 x MP1390 | 1 | 305 | 247 | 29 | 18 | 3 | 8 | 80.98 | 19.02 |
| | | | 2 | 226 | 164 | 36 | 18 | 4 | 4 | 72.57 | 27.43 |
| | | | 3 | 351 | 247 | 59 | 37 | 6 | 2 | 70.37 | 29.63 |

**Supplementary Table 8.** Analysis of recombination at arm interval *ade6-arg1* during meiosis, related to Supplementary Figure 3.

| | **Genotype** | **Strains crossed** | **No. of crosses** | **Total tested** | **Arm recombination (*ade6-arg1*)** | |
|---|---|---|---|---|---|---|
| | | | | | No. of recombinants | Recombinant Frequency (RF) (%) |
| 1 | Wild type | MP8 x MP157 | 3 | 390 | 132 | 33.8 |
| 2 | *rec11ee* | MP498 x MP499 | 3 | 389 | 149 | 38.3 |
| 3 | *rec11Δ* | MP450 x MP453 | 3 | 469 | 3 | 0.64 |
| 4 | *rec11Δ rec11ee* | MP797 x MP782 | 3 | 294 | 76 | 25.9 |
| 5 | *rec11(5A)* | MP520 x MP521 | 3 | 672 | 61 | 9.08 |
| 6 | *rec11(5A) rec11ee* | MP796 x MP883 | 3 | 411 | 122 | 29.68 |

**Supplementary Table 9.** Analysis of segregation defects at c*en1* during meiosis, related to Figures 3, 4 and Supplementary Figures 2, 3 and 4.

| | Genotype | Strains crossed | Trials | Total tested | Segregation pattern | | | | |
|---|---|---|---|---|---|---|---|---|---|
| | | | | | Normal | C-CO/RS | Meiosis I | | Meiosis II |
| | | | | | | | NDJ | PSSC | |
| 1 | Wild Type | MP208 x MP238 | 1 | 141 | 136 | 2 | 0 | 1 | 2 |
| | | | 2 | 163 | 149 | 2 | 2 | 2 | 8 |
| | | | 3 | 83 | 78 | 2 | 0 | 0 | 3 |
| | | | 4 | 133 | 127 | 2 | 0 | 0 | 4 |
| | | | 5 | 124 | 117 | 2 | 1 | 0 | 4 |
| | | | 6 | 188 | 174 | 1 | 1 | 5 | 7 |
| 2 | *psc3Δ rec8ee rec11ee* | MP368 x MP387 | 1 | 156 | 22 | 1 | 11 | 10 | 112 |
| | | | 2 | 125 | 20 | 3 | 3 | 7 | 92 |
| | | | 3 | 83 | 9 | 1 | 5 | 4 | 64 |
| 3 | *swi6Δ* | MP288 x MP366 | 1 | 128 | 57 | 1 | 2 | 4 | 64 |
| | | | 2 | 166 | 76 | 3 | 2 | 13 | 72 |
| | | | 3 | 83 | 41 | 0 | 1 | 1 | 40 |
| 4 | *sgo1Δ* | MP745 x MP746 | 1 | 178 | 6 | 6 | 14 | 9 | 143 |
| | | | 2 | 95 | 6 | 3 | 0 | 2 | 84 |
| | | | 3 | 142 | 3 | 0 | 3 | 10 | 126 |
| 5 | *ppa2Δ* | MP754 x MP770 | 1 | 141 | 114 | 3 | 3 | 3 | 18 |
| | | | 2 | 109 | 95 | 0 | 0 | 3 | 11 |
| | | | 3 | 193 | 165 | 2 | 0 | 3 | 23 |
| 6 | *chp2Δ* | MP713 x MP756 | 1 | 228 | 220 | 5 | 1 | 0 | 2 |
| | | | 2 | 247 | 222 | 6 | 4 | 4 | 11 |
| | | | 3 | 140 | 137 | 1 | 0 | 1 | 1 |
| 7 | *clr4Δ* | MP749 x MP772 | 1 | 78 | 20 | 0 | 0 | 1 | 57 |
| | | | 2 | 75 | 17 | 1 | 2 | 3 | 52 |
| | | | 3 | 62 | 18 | 0 | 1 | 1 | 42 |

| | | | | | | | | | |
|---|---|---|---|---|---|---|---|---|---|
| 7 | *psc3Δ rec8-CDee rec11-CD rec11ee* | MP720 x MP721 | 1 | 120 | 26 | 5 | 2 | 13 | 74 |
| | | | 2 | 125 | 33 | 4 | 3 | 9 | 76 |
| | | | 3 | 232 | 31 | 3 | 5 | 19 | 174 |
| 8 | *psc3Δ rec8-CDee rec11(5A)-CD rec11ee* | MP722 x MP723 | 1 | 216 | 75 | 2 | 1 | 14 | 124 |
| | | | 2 | 187 | 63 | 0 | 3 | 12 | 109 |
| | | | 3 | 244 | 43 | 4 | 1 | 22 | 174 |
| 9 | *psc3+ rec8-CDee rec11-CD rec11ee* | MP718 x MP719 | 1 | 122 | 110 | 4 | 0 | 1 | 7 |
| | | | 2 | 229 | 213 | 5 | 0 | 0 | 11 |
| | | | 3 | 57 | 49 | 1 | 1 | 1 | 5 |
| 10 | *moa1Δ* | MP1381 x MP1382 | 1 | 125 | 30 | 1 | 7 | 19 | 68 |
| | | | 2 | 292 | 58 | 3 | 10 | 48 | 173 |
| | | | 3 | 426 | 118 | 5 | 7 | 64 | 232 |

**Supplementary Table 10.** Analysis of recombination between markers flanking *cen3* and at arm interval *cen3-ade6* during meiosis, related to Figure 3.

| | **Genotype** | **Strains crossed** | **No. of crosses** | **Total tested** | ***cen3* recombination (RF)** | | | **Arm recombination (RF)** | |
|---|---|---|---|---|---|---|---|---|---|
| | | | | | Ura$^{o}$ His$^{+}$ | His$^{o}$ Ura$^{+}$ | *cen3* RF (%) | No. of recombinants | *cen3-ade6* RF (%) |
| 1 | Wild type | MP78 x MP639 | 3 | 374 | 0 | 0 | - | 61 | 16.3 |
| 2 | *sgo1*Δ | MP78 x MP639 | 6 | 683 | 0 | 1 | 0.2 | 83 | 12.2 |
| 3 | *clr4*Δ | MP78 x MP639 | 3 | 462 | 10 | 7 | 3.7 | 21 | 4.5 |

### Supplementary Discussion

Our current assay may not conclusively be able to distinguish whether the chromosomes undergoing MI NDJ experienced a C-CO or not. Even if add a third fluorophore to the other side of centromere in one of the homologs, although there will be few combinations of fluorophore colour patterns unique to a C-CO event leading to MI NDJ (Suppl. Discussion Fig. 1 below), there would still be almost similar number of segregation profiles, which will be common to MI NDJ events arising in the absence of a C-CO. This will be also true for MI PSSC events (Suppl. Discussion Fig. 2).

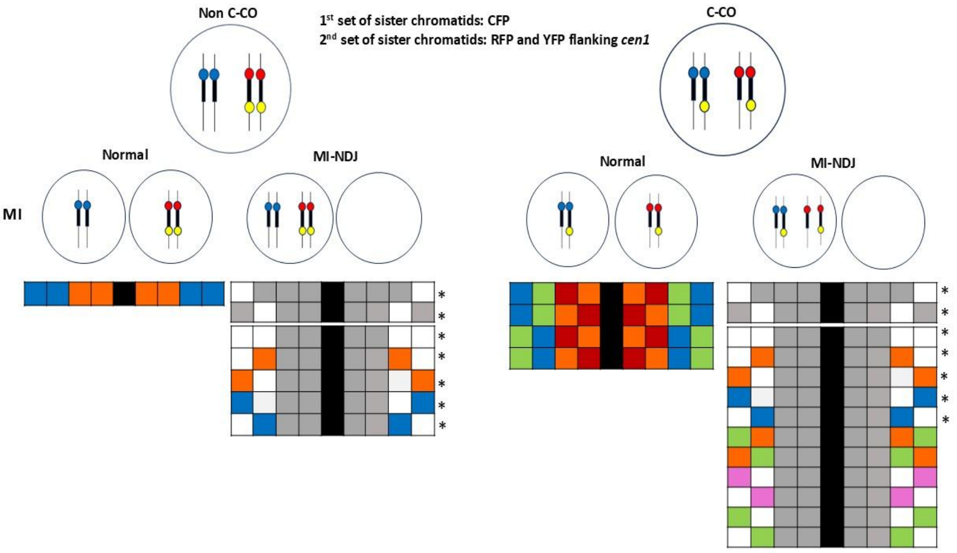


**Supplementary Discussion Figure 1.** Each of the rows depicts the combination of 4 spore colours representing the segregation profiles. The black bands are put to separate the asci that are mirror images of each other. The combinations marked with "*" represent the ones common between tetrads experiencing C-CO or not. Green signal represents a merge of CFP and YFP. Orange represents a merge of RFP and YFP; White represents a merge of CFP, RFP and YFP. Grey represents a spore with no colour.

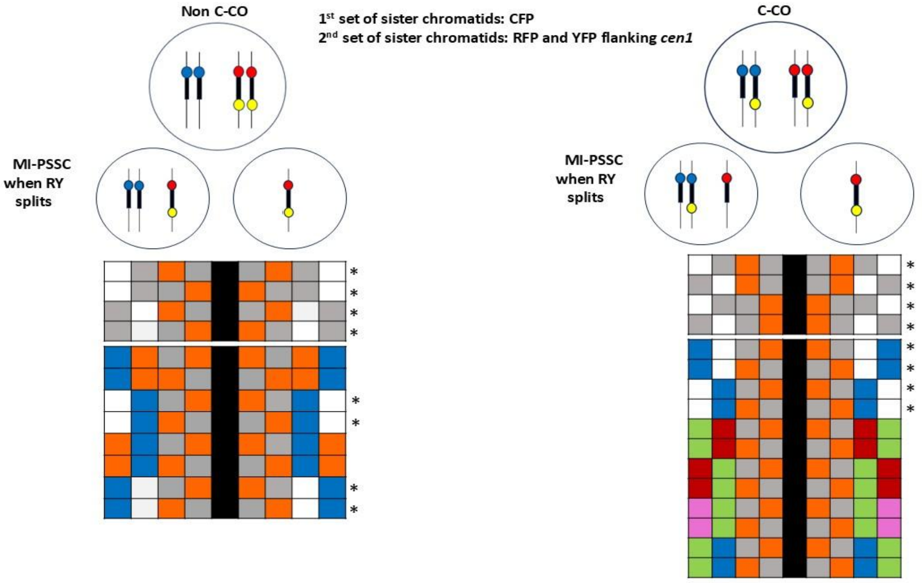


**Supplementary Discussion Figure 2.** Each of the rows depicts the combination of 4 spore colours in an ascus, representing the segregation profiles. The black bands separate the asci which are mirror images of each other. The combinations marked with "*" represent the ones common in C-CO and non

C-COs. Green signal represents a merge of CFP and YFP. Orange represents a merge of RFP and YFP; White represents a merge of CFP, RFP and YFP. Grey represents a spore with no colour.

As can be clearly seen in both the cases of MI defects, the errors arising due to random homolog disjunction will be a subset of those arising due to C-COs, making it impossible to distinguish between the two. The list of patterns shown here are not exhaustive, but clearly suggest that using the three-fluorophore system will not unequivocally distinguish between MI errors arising from C-COs and those without.

Therefore, in our study we attempted to compensate for these caveats by keeping sufficient controls to be able to infer with the best certainty that these MI segregation defects are occurring due to C-COs. Of particular note is the *rec10(mut)-CD rec10ee* control, where the MI errors are significantly lower than *rec10-CD rec10ee,* which we think indicates strongly that the errors we are seeing should be due to an increase in C-COs during meiosis. We additionally overexpressed *rec10* in these backgrounds to compensate for any deficiencies that may occur due to Rec10-associated functions in linear element formation. In the absence of an alternate hypothesis, we think that the MI errors seen in the presence of Rec10-CD *rec10ee* can be explained due to presence of C-COs. Furthermore, known cohesion and recombination mutants used by us, faithfully recapitulate previously reported patterns of mis-segregation suggesting that our assay is capturing the defect phenotypes faithfully and that these are different from the ones we think are occurring due to C-COs.
